# Shaped by shape: costs and topology drive host-parasite evolution in spatial networks

**DOI:** 10.64898/2026.11.18.767619

**Authors:** Saismit Naik, Jhelam N. Deshpande, Iago Bonnici, Oliver Kaltz, Emanuel A. Fronhofer

**Author notes:** **Correspondence Details**, Emanuel A. Fronhofer, Institut Des Sciences De L’Evolution De Montpellier, France.

## Abstract

How disease spreads across space is well studied, but how the topology of spatial networks shapes host-parasite evolution is less understood. Host-parasite interactions unfold across spatial networks in which populations differ in the number of neighbours, yet most theory has focused on regular networks such as grids. Here, we embed a multilocus gene-for-gene infection model into a metapopulation to investigate host-parasite coevolution across empirically inspired and synthetic spatial networks.

We find that variation in the number of neighbours generates spatial variation in local host density and parasite prevalence, creating a network-wide mosaic of ecological conditions for selection. Greater variation in the number of neighbours, or higher degree heterogeneity, promotes the evolution of host resistance. The same degree heterogeneity leads to reduced parasite infectivity in well-connected patches versus higher infectivity in patches with fewer neighbours. Both outcomes arise because populations with fewer neighbours experience net host outflux and reduced prevalence, whereas populations with more neighbours accumulate hosts and experience different selective pressures. Dispersal both drives and homogenises host resistance across the network, while maintaining spatial structure in parasite infectivity. We here identify a topology-epidemiology-evolution feedback impacting coevolution even when populations experience identical environmental conditions. Across network types, coevolutionary outcomes are driven by the network’s degree heterogeneity, which can be quantified using the Herfindahl-Hirschman Index in real landscapes. Our results provide testable predictions for how changes in network topology can shape the evolution of host resistance and parasite infectivity.

**Significance Statement:** Anthropogenic disturbances are causing unprecedented rates of change to connectivity of habitats, including our own. We show that just the network topology of a metapopulation can predictably reshape host-parasite coevolution. In a metapopulation, variation in the number of neighbours among otherwise equivalent populations generates differences in population density through asymmetric dispersal, producing higher host densities in highly connected populations and lower densities in sparsely connected ones. The asymmetry in densities in turn produces a spatial mosaic of epidemiology that drives selection: sparsely connected populations favour host resistance, whereas high-density populations favour reduced parasite infectivity. By linking these mechanisms to measurable network indices, our results provide testable predictions for how changes in connectivity shape host resistance and parasite infectivity.

## 1 Introduction

Hosts and parasites are locked in an evolutionary arms race (Anderson and May, 1982): hosts evolve resistance, while parasites evolve the infectivity needed to overcome it. Because parasites depend on encountering susceptible hosts, this arms race unfolds not only through time but also across space: the pattern of contacts among host populations determines where parasites persist, how widely they spread, and which host and parasite genotypes are favoured. Although epidemiology has shown that contact-network structure can profoundly alter disease spread (Colizza and Vespignani, 2007; Volz et al., 2011; Tian et al., 2020), we still lack a mechanistic understanding of how heterogeneous spatial networks shapes the coevolution of host resistance and parasite infectivity. Indeed, a review of host-parasite coevolutionary models found that only 14.5% (27 of 185 studies) considered space in any form (Buckingham and Ashby, 2022).

In theoretical studies of host-parasite coevolution that have considered spatial structure, space is often represented using regular lattices (Boots and Sasaki, 1999; Gandon, 2002; Ashby et al., 2014) or two-patch systems (Peters et al., 2023). These approaches have provided important insights into how spatial structure and local interactions shape coevolution, but they do not capture the heterogeneity in connectivity that is characteristic of empirical spatial metapopulations, where patches differ in the number and arrangement of neighbours. Such variation can generate spatial structure in local population densities, influence metapopulation dynamics (Kininmonth et al., 2010; Fronhofer and Altermatt, 2017), as well as generate phenotypic differentiation across spatial networks (Boussange and Pellissier, 2022). In host-parasite systems, these demographic differences can alter local infection prevalence and transmission, thereby changing the conditions under which selection on host resistance and parasite infectivity operates. The evolutionary consequences of these local conditions may also depend on the costs of resistance and infectivity, which determine the relative benefits of evolving each trait.

Empirical work across a variety of systems has shown that spatial arrangement and habitat fragmentation can influence host-parasite dynamics and evolution (Guivier et al., 2011; Jousimo et al., 2014; Carlsson-Granér and Thrall, 2015; Parratt et al., 2016; Testa et al., 2019; Höckerstedt et al., 2022). For example, landscape-genetic studies of bank voles and Puumala hantavirus have shown that viral prevalence depends not only on local density but also on metapopulation connectivity and effective population size (Guivier et al., 2011). In the *Plantago lanceolata*-*Podosphaera plantaginis* system, well-connected host populations experienced lower pathogen colonisation, higher pathogen extinction rates and higher resistance than isolated populations, consistent with an influence of connectivity and gene flow on host-pathogen dynamics (Jousimo et al., 2014; Höckerstedt et al., 2022). Recent theoretical studies focusing on parasite evolution have shown that spatial variation in local environments can alter parasite virulence (Sato et al., 2024), while network topology can influence virulence evolution in river-like and terrestrial-like networks (Deshpande et al., 2025a,b). However, it remains unclear how network topology generates spatial variation in the ecological and epidemiological conditions shaping host resistance and parasite infectivity. It is also unclear how these local evolutionary responses translate into network-wide coevolutionary outcomes.

Here, we investigate how spatial network topology shapes host-parasite coevolution through its effects on local ecological and epidemiological conditions. Our central question is: How does network topology shape the coevolution of host resistance and parasite infectivity? We develop an individual-based, eco-evolutionary susceptible-infected (SI) model (Kermack and McKendrick, 1927) incorporating multilocus gene-for-gene (GFG) genetics (Flor, 1956; Sasaki, 2000), in which host resistance and parasite infectivity evolve across spatially structured host populations. We compare empirically inspired network types, including river-like optimal channel networks (Carraro et al., 2020) and terrestrial-like random geometric graphs (Gilarranz and Bascompte, 2012), with homogeneous controls and synthetic networks spanning gradients of degree heterogeneity and assortativity.

We show that network topology generates spatial variation in host density and parasite prevalence, creating a mosaic of local epidemiological conditions that shape the evolution of host resistance and parasite infectivity. Importantly, the local epidemiology is not necessarily reflected in equivalent spatial patterns of evolved traits: dispersal can homogenise host resistance across the landscape, whereas parasite infectivity retains a stronger spatial structure. By identifying how degree heterogeneity and assortativity jointly shape these ecological and evolutionary processes, we provide a mechanistic framework for understanding how spatial network topology influences host-parasite coevolution. Our results highlight degree heterogeneity, quantified using the Herfindahl-Hirschman Index, and degree assortativity, quantified using Spearman’s rank correlation (*ρ*), as measurable network properties associated with coevolutionary outcomes.

## 2 Result and Discussion

### 2.1 River-like networks select for increased resistance and infectivity

Coevolutionary dynamics in our multilocus gene-for-gene (GFG) model are strongly shaped by the costs of resistance and infectivity, as well as by its characteristic asymmetric infection genetics (Fig. 1; Section 4.1). Host and parasite genotypes are binary sequences of length*n*= 4, yielding a four-locus, two-allele model. A host allele encoded as “1” confers resistance at the corresponding locus, whereas a “0” denotes susceptibility. Conversely, a parasite allele encoded as “1” confers infectivity at that locus, whereas a “0” denotes the absence of infectivity. A host resistance allele protects against infection only if the parasite lacks the matching infectivity allele. Thus, increasing the number of host resistance alleles reduces the probability of infection, whereas increasing the number of parasite infectivity alleles increases it (Section 4.1).

**Figure 1.**
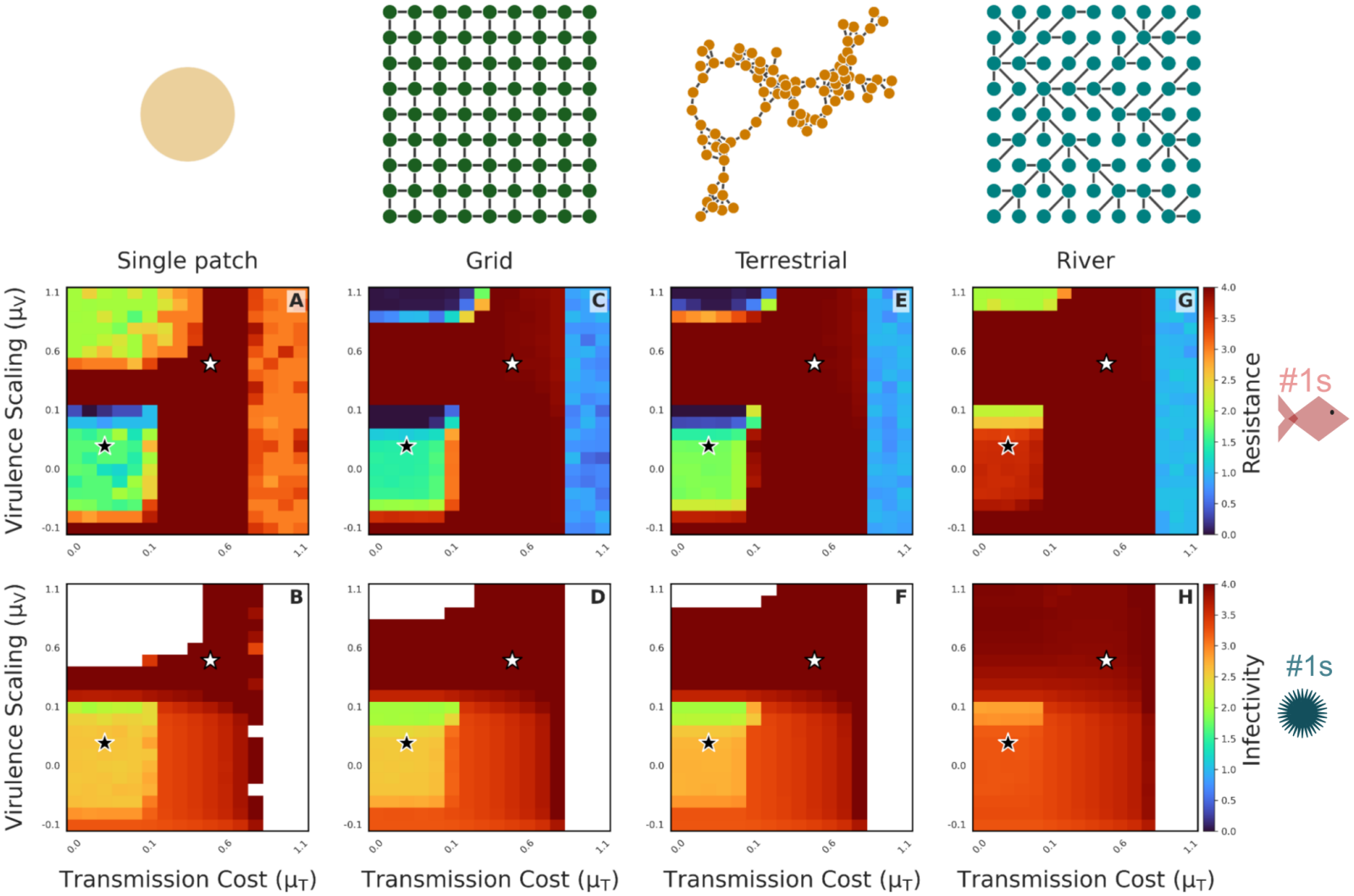
Effects of transmission cost scaling (*µ*_*T*_ ; x-axis) and virulence scaling (*µ*_*V*_ ; y-axis) on host-parasite coevolution across four network types at low resistance cost (*µ*_*R*_ = 0.000256). Each pair of panels corresponds to a specific spatial structure: (A, B) a single, well-mixed patch; (C, D) a homogeneous grid (torus); (E, F) terrestrial-like networks modelled as Random Geometric Graphs (RGGs); and (G, H) river-like networks modelled as Optimal Channel Networks (OCNs). The top row (A, C, E, G) shows evolved host resistance, quantified as the median number of resistance alleles (‘1’s) per patch across the host population, while the bottom row (B, D, F, H) shows evolved parasite infectivity, quantified as the median number of infectivity alleles (‘1’s) per patch. Colour intensity ranges from blue (low) to yellow (high), with warmer colours indicating greater evolutionary investment in resistance or infectivity. The dispersal rate is fixed to*d*= 0.5. The black star indicate the specific parameter combinations selected for detailed case studies presented in Figs. 2. Refer to Table S1 for fixed parameters. The white star indicates the parameter combination used for Fig. S1. An exploration of the impact of space with other parameters can be found in Fig. S2, S3 and S4.

In the absence of spatial structure (Fig. 1A, B), our model recapitulates classical GFG predictions (Fenton and Brockhurst, 2007; Ashby et al., 2019). Increasing transmission cost (*µ*_*T*_) eventually eliminates infectivity alleles. In our finite, stochastic model, this occurs through parasite extinction rather than the deterministic loss of infectivity alleles in infinite populations (Fenton and Brockhurst, 2007). Similarly, increasing the cost of resistance (*µ*_*H*_) eventually eliminates resistance alleles (Figs. S7, S8). Because variation in*µ* _*H*_ produces largely binary outcomes, resistance either evolves or it does not (Figs. S5 and S6). We here focus on low*µ* _*H*_ , where resistance can evolve, and vary the two parasite-associated costs: transmission cost (*µ*_*T*_) and virulence (*µ*_*V*_) (Gandon et al., 2002).

Relative to a single well-mixed population, homogeneous landscapes (Fig. 1 C, D for grids and Fig. 1 E, F for RGGs), produce no striking differences apart from an expected “metapopulation effect” (less extinctions in Fig. 1 D and F; see results with respect to*µ* _*H*_ in Fig. S8). Parasites that would otherwise go extinct under high transmission or virulence costs in a single-patch can persist across space (Deshpande et al. 2025a, S14; Tadiri et al. 2018).

However, variation in number of neighbours of patches, that is, degree heterogeneity, as in OCNs (Fig. 1 G, H) leads to interesting, diverging results: OCNs exhibit a pronounced increase in host resistance and parasite infectivity at low costs. Correlations between the coevolutionary results of infectivity alleles and resistance alleles across the parameter space of Fig. 1 can be found in Fig. S9.

### 2.2 Degree heterogeneity promotes host resistance and generates spatial structure in parasite infectivity

To investigate the mechanisms underlying the network-topology-driven differences in coevolution described in Section 2.1, we examine the evolution of host resistance and parasite infectivity across dispersal rates under low virulence (*µ*_*V*_ = 0.0001) and low transmission costs (*µ*_*T*_ = 0.0001). This parameter combination is indicated by the black star in Fig. 1. Our results in this section apply to the parameter region in which host resistance has not reached its upper bound (i.e., the regions that are not red in Fig. 1A).

In OCNs, evolved resistance increases rapidly with dispersal before reaching a maximum, whereas resistance increases more gradually in RGGs and changes little in regular networks (Fig. 2A, B). Thus, the effect of dispersal on resistance evolution depends strongly on network topology, particularly on the presence of degree heterogeneity.

**Figure 2.**
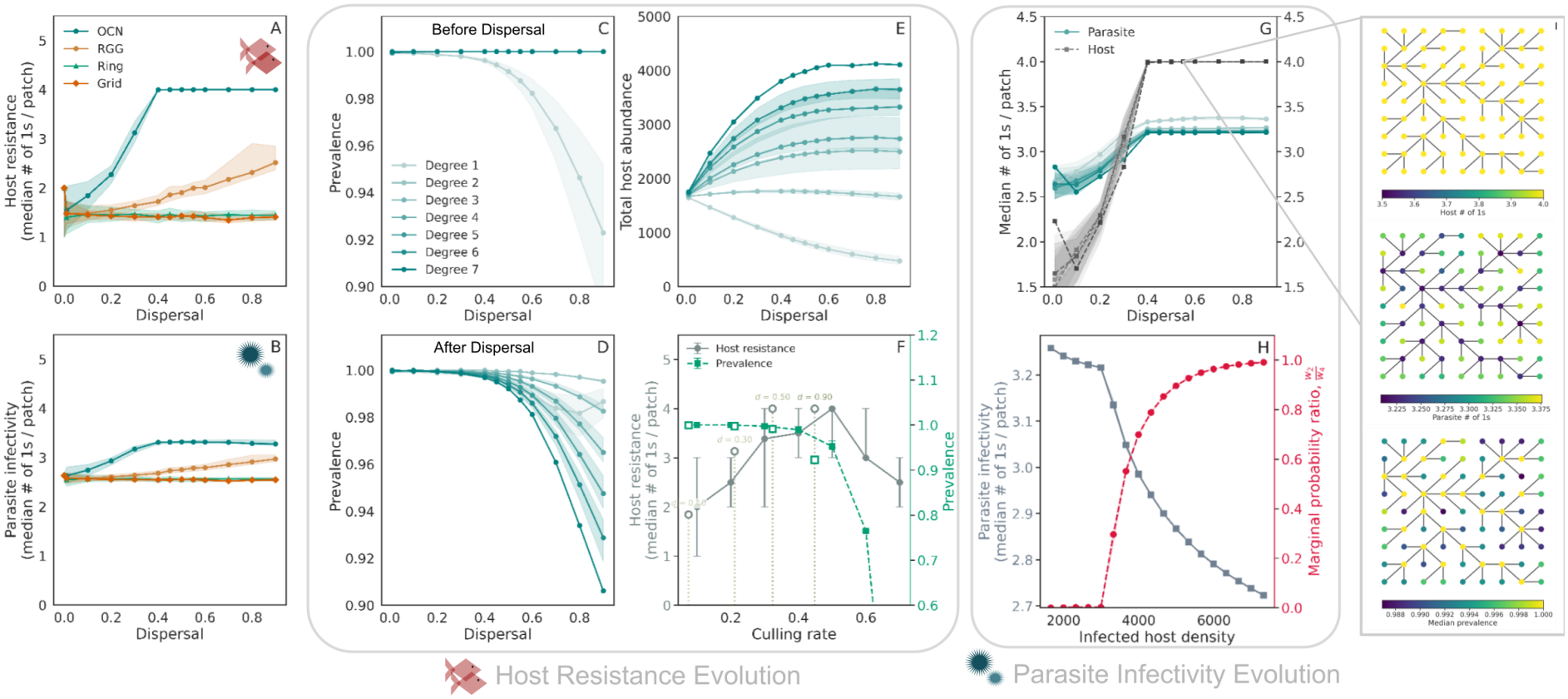
Impact of dispersal and network-type on coevolutionary outcomes when costs are low (virulence scaling*µ* _*V*_ = 0.0001, transmission cost scaling*µ* _*T*_ = 0.0001, resistance cost scaling*µ* _*R*_ = 0.000256, black star in Fig. 1). Panel A) Host resistance (median number of “1” alleles in the host per patch) as a function of dispersal. Results for the OCNs are shown in teal, the RGG in orange, the ring and grid networks in vermilion and sea green respectively. Shaded regions always give the interquartile range across patches. Parasite infectivity (median number of “1” alleles in the parasite) per patch as a function of dispersal. Median prevalence before dispersal per patch grouped by their degrees in the OCNs. D) Median prevalence after dispersal per patch grouped by their degrees in the OCN. E) Median host abundance after dispersal per patch grouped by their degrees in the OCN. F) Simulation results from a single-patch with host resistance on the y-axis on the left, prevalence in the patch on the right y-axis and increasing culling rate on the x-axis. The vertical lines are drawn at median effective culling experienced through net host outflux in the single-degree patches of the OCNs at dispersal rates 0.1,0.3,0.5,0.9. The unfilled markers correspond to the median prevalence and resistance experienced the single-degree patches. G) Host resistance/parasite infectivity per patch as a function of dispersal, grouped by patch degree in the OCN. Teal lines show parasite infectivity and grey lines (light to dark with increasing degree) show host resistance. H) Parasite infectivity (solid grey, left axis) and the median ratio of marginal probability of parasite genotypes with 2 “1” or infectivity alleles (*w*_2_) to marginal probability of parasite genotypes with 4 “1” alleles (*w*_4_) as defined by Eqn. S2 (dashed crimson, right axis) as functions of equilibrium infected host densities in a single-patch (also see *w* _*k*_ plotted as functions of local infected densities in Fig. S10). I) Spatial distribution of host resistance, parasite infectivity, and prevalence across a single OCN realisation (measured before dispersal). Patch colour indicates the weighted number of “1” alleles per host patch, parasite patch, and the median prevalence. Refer to Table S1 for fixed parameters. Sensitivity analyses for panels A and B can be found in Fig. S11.

We show that the increase in resistance in degree-heterogeneous networks, such as OCNs and less strongly in RGGs, is driven by spatial variation in infection prevalence. Degree heterogeneity generates net host outflux from low-degree patches, maintaining lower local host density than in the absence of dispersal (Fig. S14). This reduction in local host density lowers local infection prevalence (Fig. 2C; Supplementary Information S4.7). After transmission and before dispersal (Fig. S13), prevalence declines with increasing dispersal in single-degree patches (Fig. 2C; see Fig. S12 for spatial patterns before and after dispersal), but remains high in higher-degree patches. High prevalence can weaken selection for resistance because resistance provides little benefit when most hosts are already infected (Walsman et al., 2023). After dispersal, net host outflux from low-degree patches and net host influx into high-degree patches (Fig. S14) reduce prevalence in high-degree patches (Fig. 2D) but the cycle repeats in the next transmission phase, when remaining susceptible hosts become infected.

To confirm that net host outflux from low-degree patches is responsible for lower prevalence and stronger selection for resistance, we mimicked net host outflux by culling a fraction of the population in a single-patch simulation. Increasing the culling rate reduced equilibrium prevalence and increased resistance (Fig. 2F), reproducing the qualitative pattern observed in single-degree patches. Analytically, increasing culling always reduces equilibrium prevalence in our model, such that d*p*^*\**^*/*d*c <*0 (Supplementary Information S4.7). Moreover, equilibrium resistance and prevalence in single-degree patches of OCNs and RGGs follow the qualitative trends expected from their respective net host outflux (vertical lines and unfilled markers in Fig. 2F and Fig. S15F). Single-degree patches show higher resistance and lower prevalence than the corresponding culling treatment because they continue to receive resistant host immigrants and parasites from connected high-degree patches. Together, these results show that degree-dependent net host outflux reduces local density and prevalence, creating conditions that favour resistance evolution.

Resistance selected under these low-prevalence conditions can then spread through the network via dispersal, contributing to the network-wide increase in resistance observed in OCNs. The strength of this effect depends on the degree distribution. RGGs show the same qualitative pattern in single-degree patches (Fig. S15; Fig. S12), but contain relatively few single-degree patches (Fig. S16) and have a more dispersed degree distribution than OCNs. Consequently, RGGs generate less pronounced variation in local host abundance and weaker selection for resistance (Fig. S15E).

Degree heterogeneity also generates spatial structure in parasite infectivity through a distinct, density-dependent transmission mechanism. Parasite infectivity declines with patch degree: Single-degree patches harbour parasites with higher infectivity, whereas high-degree patches harbour the parasites with lesser infectivity (Fig. 2G; see Fig. 2I for a single-replicate example). Single-degree patches experience net host outflux, whereas high-degree patches experience net host influx and maintain higher equilibrium host densities than in the absence of dispersal (Fig. S14). Higher host densities also imply higher infected host densities. At high infected host density, both the probability of infection (Eqn. 6) and the marginal probability of infection by a given parasite genotype (Eqn. S2; Fig. S10) approach one. Additional infectivity alleles therefore provide diminishing marginal benefits to a parasite genotype (Fig. S10). Because parasites with the maximum number of infectivity alleles have the highest per-parasite transmission rate (Eqn. 6), their infection probability saturates fastest with increasing local infected host density (Fig. S10). Lower-infectivity genotypes saturate more slowly and therefore gain a relative advantage as infected host density increases.

Parasites in high-degree patches therefore evolve lower infectivity because they experience greater transmission probability saturation: The advantage of higher number of infectivity alleles in parasites declines as infected host density increases. To recapitulate this pattern, we increased the host intrinsic growth rate (*λ*_0_) in a single-patch simulation. As the number of infected individuals rises, the marginal infection probability of high-infectivity genotypes saturates, and the relative advantage of lower-infectivity genotypes increases (see*w* _2_*/w*_4_ in Fig. 2H; Supplementary Information S4.3).

This topology-epidemiology-evolution feedback, which reduces parasite infectivity in high-degree patches, also has important consequences in the parameter regime where host resistance evolves to its maximum (white star in Fig. 1;*µ* _*V*_ = 0.5,*µ* _*T*_ = 0.6). In this parameter regime, selection maximises host resistance but introduction of spatial structure reduces parasite infectivity at the network scale in OCNs (Fig. S1B). The full analysis of this scenario is presented in Supplementary Information S1.1. The degree-driven spatial structure in infected host density remains the same (Fig. S1C), causing density-dependent transmission to saturate in high-degree patches. This weakens the marginal benefit of additional infectivity and favours lower-infectivity genotypes (Fig. S1E). Consequently, high-degree patches maintain lower parasite infectivity, whereas low-degree patches maintain maximum parasite infectivity because they experience lower transmission saturation and therefore retain stronger selection for high infectivity.

As degree-driven spatial variation in infected host density increases, high-degree patches therefore reduce network-wide parasite infectivity.

Together, these results show that degree heterogeneity generates spatially heterogeneous epidemiological conditions, with distinct consequences for host resistance and parasite infectivity coevolution. Resistance evolves in response to the epidemiological consequences of net host outflux from low-degree patches, but is subsequently redistributed through the network by dispersal, reducing spatial structure in resistance. Parasite infectivity, by contrast, responds more directly to local infected host density, generating pronounced spatial variation in infectivity across patches.

### 2.3 Degree heterogeneity and assortativity drive coevolution

Independent of the exact network topology, degree heterogeneity and degree assortativity jointly shape local population density and coevolutionary outcomes (Fig. 3, Fig. S20). Degree heterogeneity measures variation among patches in their number of neighbours, whereas degree assortativity measures whether patches with similar numbers of neighbours tend to connect to one another. Changing these topological properties can concentrate dispersal in well-connected patches or diffuse the distribution of population densities. We generated networks spanning a two-dimensional gradient of these properties (see Supplementary Information S4.6 for details). A sub-sample of the networks generated during the exploration along with other network metrics can be found in Fig. S26. We measure degree heterogeneity of a network using the Herfindahl-Hirschman Index (HHI) that quantifies concentration of node connections and we measure assortativity using the Spearmans rank correlation coefficient (*ρ*) (Litvak and Van Der Hofstad, 2013).

**Figure 3.**
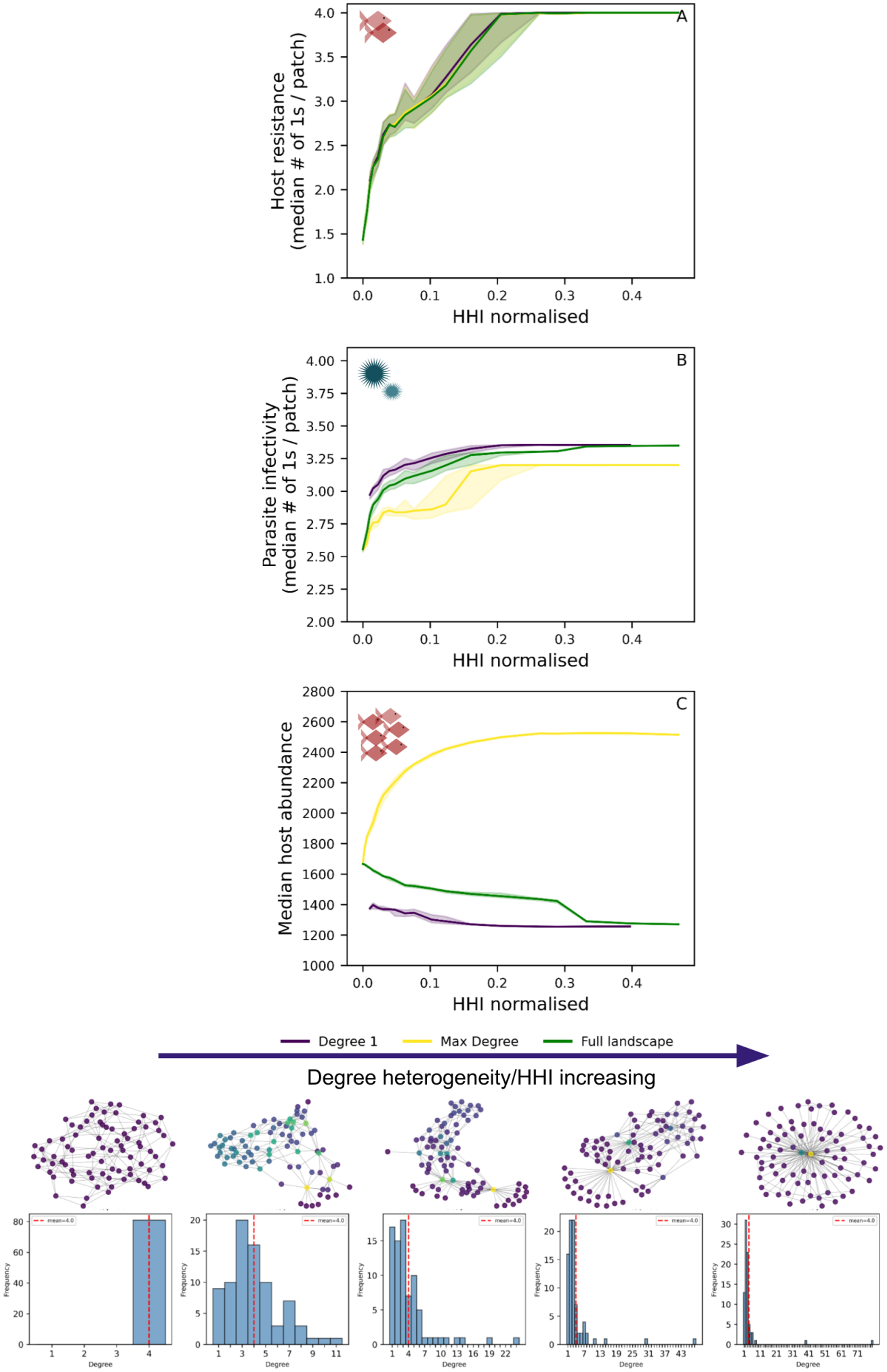
Coevolutionary outcomes and host abundance as a function of degree heterogeneity. The x-axis shows degree heterogeneity measured as the normalised Herfindahl-Hirschman index (*HHI* _norm_; Supplementary Information S4.5.1) across networks generated using edge-rewiring algorithms on a regular 4-connected graph with *n*= 81 patches (see Supplementary Information S4.6) . For each HHI level median and interquartile range (IQR) are computed across networks with different assortativity (quantified by Spearman’s rank correlation coefficient*ρ*; Supplementary Information S4.5.2). Three patch degree subsets are shown: violet-patches with degree 1; yellow-patches with the maximum degree found for each *HHI* _*norm*_ and Spearman’s *ρ*) combination; green-the full network (all patches with degree*≥*1). The three panels correspond to low virulence scaling (*µ*_*V*_ = 0.0001) and low transmission cost (*µ*_*T*_ = 0.0001) as in Fig. 2 and S15 and marked by the black star in Fig.1. A) Median host resistance (number of “1” alleles per patch). B) Median parasite infectivity (number of “1” alleles per patch). C) Median host abundance per patch. Dispersal is fixed at *d*= 0.3. All other fixed parameters are listed in Table S1. At the bottom, we show a visualisation of networks and corresponding degree-distribution across HHI values.

Increasing degree heterogeneity or HHI generates stronger spatial structure in local population density (between high and low-degree patches; Fig. 3C, Fig. S20C). When virulence and transmission costs are low (*µ*_*V*_ = 0.0001, *µ* _*T*_ = 0.0001), greater degree heterogeneity is associated with higher network-wide host resistance (Fig. 3 A, see with dispersal, *d*= 0.5 in Fig. S19). This pattern is consistent with results reported in Fig. 2: Greater degree heterogeneity generates stronger differences in local density and prevalence, particularly in sparsely connected patches, thereby strengthening selection for resistance. The trends remain robust to different average degrees (Fig. S23 with a 2-regular graph, Fig. S22 with a 8-regular graph) as well as different numbers of patches (Fig. S24 with 40 patches, Fig. S25 with 160 patches).

When virulence and transmission costs are high (*µ*_*V*_ = 0.5,*µ* _*T*_ = 0.6) and host resistance is maximised, greater degree heterogeneity is associated with lower parasite infectivity (Fig. S20 A-B, see with dispersal, *d*= 0.5 in Fig. S21). This result is consistent with results reported in Fig. S1, in which density heterogeneity generated by OCNs favours reduced infectivity in high-degree patches. Thus, the evolutionary effect of degree heterogeneity depends on which host or parasite trait remains able to evolve.

Degree assortativity further modifies these effects: At a given level of degree heterogeneity, increasing assortativity, that is, increasing the tendency for populations with similar numbers of neighbours to connect, is associated with lower host resistance (Fig. S27 A, Fig. S28 A) and higher parasite infectivity (Fig. S29 B, Fig. S30 B) at the network level. These evolutionary changes coincide with differences in the distribution of local population density across the network (Fig. S27 C, Fig. S28 C, Fig. S29 C, Fig. S30 C). Thus, the arrangement of connections matters in addition to the degree distribution: Even networks with similar degree heterogeneity can generate different coevolutionary outcomes depending on whether high-degree populations connect primarily to one another or to low-degree population Together, these results show that degree heterogeneity and degree assortativity jointly shape the spatial distribution of local ecological conditions and thereby predict the direction and magnitude of host-parasite coevolution. These measurable network properties provide a tractable framework for forecasting coevolutionary outcomes in natural spatially structured populations.

## 3 General Discussion

We show that degree heterogeneity, a ubiquitous feature of real-world spatial networks, can act as a crucial driver of host-parasite coevolution, with costs modulating the strength and direction of these effects. Dispersal on degree-heterogenous networks, that is, networks with variation in degrees, creates net host outflux from low-degree patches and net host influx into high-degree patches (Fronhofer and Altermatt, 2017), reducing host density and prevalence in the former while increasing host abundance in the latter (Figs. 2E, S14B). Reduced density lowers equilibrium prevalence (Fig. 2F; Supplementary Information S4.7), creating conditions under which resistance becomes beneficial. Conversely, when host resistance evolves to its upper bound in the absence of dispersal, spatial structure leads to elevated density in high-degree patches, shifting selection towards reduced parasite infectivity (Fig. S1 C, D). Our synthetic networks indicate that these effects are governed by degree heterogeneity and assortativity, rather than by idiosyncracies of network type (Fig. 3).

Our results relate to geographic mosaic theory of coevolution (Thompson, 2019), which posits that environmental heterogeneity in space generates mosaics of selection pressures. Our framework shows that heterogeneous network topology can generate a mosaic of ecological conditions for selection even when all populations experience identical environmental conditions but since we always have reciprocal selection we do not generate hot and cold-spots of coevolution. Low-density, low-degree patches experience conditions that favour resistance, whereas high-density patches can favour reduced parasite infectivity. Crucially, this selection mosaic does not always produce a matching trait mosaic: dispersal redistributes resistance alleles across the network, so that host resistance can increase network-wide without strong spatial differentiation. Parasite infectivity, by contrast, retains a stronger spatial signature because its evolution depends on the number of infected individuals locally (Fig. 3). This aligns with eco-evolutionary models showing that spatial-graph topology alone can generate phenotypic differentiation (Boussange and Pellissier, 2022). Network topology should therefore be considered a fundamental driver of coevolutionary mosaics. Our result also complement recent theory on parasite evolution suggesting that metapopulation heterogeneities can increase pathogen virulence and infectiousness (Sato et al., 2024). Our coevolutionary model is broadly consistent with the parasite-evolution result of Sato et al. (2024), but it also shows that network topology can generate spatial structure in parasite infectivity and can select for reduced infectivity when host resistance is constrained.

Our results resonate with empirical hostparasite systems: In the *Viscaria-Microbotryum* system, continuously or patchily connected host populations were more resistant and harboured more infective pathogen strains than highly isolated populations (Carlsson-Granér and Thrall, 2015), consistent with the joint escalation of resistance and infectivity across our metapopulation in the low-cost regime (Fig. 2A,B). In the *Plantago lanceolata*-*Podosphaera plantaginis* system, highly connected populations within a metapopulation were more resistant than less-connected populations (Jousimo et al., 2014; Höckerstedt et al., 2022). This pattern is broadly consistent with our finding that greater dispersal can promote resistance at the landscape level, but differs from the weak spatial structure in resistance predicted by our model. In our model, dispersal redistributes resistance across the network, reducing spatial structure in host resistance, whereas the empirical system may retain stronger local differences in resistance. This difference may reflect more complex dispersal patterns in the empirical system, which is not characterised by explicit co-dispersal of parasites with hosts but may involve correlated movement of wind-dispersed parasites and pollen. Further work is required to understand how the topology-epidemiology-evolution feedback we identify, arises in biologically realistic settings.

Our framework also addresses disease-prevalence patterns: High-degree patches maintain higher prevalence because net host influx increases host abundance and therefore the density of infected individuals (Fig. 2D). This is consistent with landscape-genetic studies of bank voles and Puumala hantavirus, in which prevalence depends on local density as well as metapopulation connectivity and effective population size (Guivier et al., 2011). Our culling experiment provides a mechanistic link: Reducing density in a single-patch lowers prevalence and favours resistance, supporting the density mechanism inferred from degree-dependent net dispersal flux in the network (Fig. 2F; Supplementary Information S4.7). Thus, network topology can reshape local prevalence through its effects on host redistribution, population density and the evolution of parasite traits.

Although our model assumes discrete generations and obligate co-dispersal, these assumptions are consistent with a broad class of host-parasite systems with strongly seasonal or pulsed transmission, including salmonid diseases such as Atlantic salmon-*Gyrodactylus salaris*and Viral Haemorrhagic Septicaemia (Appleby and Mo, 1997; Garver and Hawley, 2021), and the *Podosphaera* and *Microbotryum* systems discussed above. Several assumptions nevertheless define the scope of our predictions: Our gene-for-gene framework yields stable spatial parasite polymorphisms and network-wide resistance, rather than the fine-scale local adaptation often generated by matching-allele models (Gandon, 2002; Kim and Ashby, 2026). Because natural systems may lie along a continuum between these genetic architectures (Agrawal and Lively, 2003; Buckingham and Ashby, 2022), testing how alternative infection genetics interact with network topology is an important future research avenue. We also assume unbiased dispersal with respect to infection status, undirected and unweighted networks, and a single parasite species. Directionality, infection-dependent movement, patch extinction, environmental variation, and multiple pathogen strains could all alter the density gradients and selection mosaics we describe (Lion and Boots, 2010; Fron-hofer and Altermatt, 2017). Incorporating these processes will clarify when network topology generates, maintains, or erodes coevolutionary diversity.

More broadly, anthropogenic change is altering the topology of spatial networks through habitat loss, fragmentation, land-use conversion, and climate-driven range shifts (Bullock et al., 2018). Human disturbance may produce more heterogeneous networks in which a few populations have many connections and therefore higher degree heterogeneity (Bullock et al., 2018). Our framework predicts that such changes will shift host-parasite coevolutionary trajectories, potentially selecting for increased infectivity depending on the underlying cost architecture. Because degree heterogeneity and assortativity are measurable properties of real-world spatial networks, they may help identify where coevolutionary responses are most likely and inform disease management in rapidly changing environments.

## 4 Model description

### 4.1 Infection genetics and costs

Genetic specificity follows a multilocus gene-for-gene (GFG) framework (Sasaki et al., 2002). Host and parasite genotypes are binary sequences of length *n* that determine infectivity, transmission, and fitness costs. Coevolution occurs through mutation at each locus, and we track the densities of host and parasite genotypes across space and time.

Genetic specificity follows a multilocus gene-for-gene (GFG) framework (Sasaki et al., 2002). Hosts and parasites are haploid and carry binary genotypes at *n*= 4 loci, yielding 2 ^4^ = 16 possible genotypes per species. Each host *i* has genotype 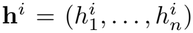, where 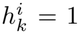 denotes a resistance allele and 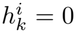 its absence. Each parasite *j* has genotype 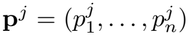, where 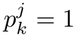 denotes an infectivity allele. A host resistance allele provides protection only when the corresponding parasite infectivity allele is absent. Thus, the number of effective resistance loci is

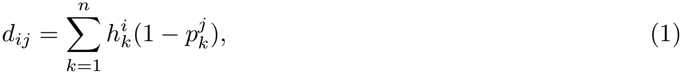

and the infectivity of parasite *j* on host *i* is

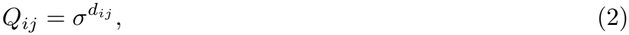

where*σ*is the reduction in infectivity caused by each effective resistance locus (Sasaki, 2000; Sasaki et al., 2002; Ashby et al., 2014).

Host resistance reduces fecundity by

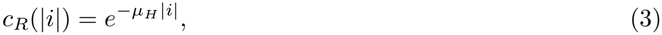

Where 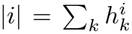 is the number of resistance alleles and*µ* _*H*_ scales the cost of resistance (Sasaki, 2000; Ashby et al., 2014). Infected hosts experience an additional fecundity reduction due to parasite virulence. It is the cost of expressing those alleles, whereby greater exploitation reduces host fecundity and infectious period indirectly (Anderson and May, 1982; Alizon and van Baalen, 2005),

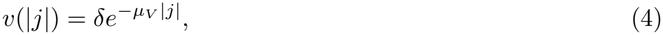

Where 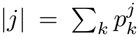 is the number of parasite infectivity alleles, *δ* is the baseline fecundity of an infected relative to a susceptible host, and *µ* _*V*_ scales virulence. Parasites with more infectivity alleles therefore impose greater virulence but can pay a transmission cost. It represents the classical GFG cost of carrying infectivity alleles as cost of generalism, whereby broader host range reduces transmission efficiency (Sasaki, 2000; Fenton and Brockhurst, 2007; Ashby et al., 2014),

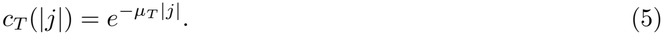

For host *i* and parasite *j*, effective transmission is therefore

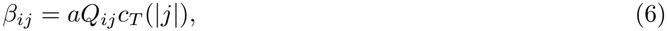

Where *a* is search efficiency (Deshpande et al., 2025a). This formulation separates the cost of carrying infectivity alleles from the cost of expressing them through virulence, allowing us to examine their joint effects on coevolution (Anderson and May, 1982; Gandon et al., 2002; Alizon and van Baalen, 2005).

### 4.2 Life cycle

At the beginning of each generation, hosts disperse with probability *d* from their natal patch to a randomly chosen connected patch. Parasites only disperse with infected hosts (co-dispesral; (North and Godfray, 2017)). Dispersal is governed by the binary adjacency matrix **D**, where *D* _*pq*_ = 1 if patches *p* and *q* are connected. A schematic overview of the simulation cycle, from dispersal through transmission to parasite mutation, is shown in Fig. S13.

After dispersal, hosts reproduce locally according to Beverton-Holt density dependence (Beverton et al., 1957). Hosts and parasites are haploid and reproduce asexually. The system has non-overlapping generations, and each generation consists of dispersal, reproduction, mutation, and transmission. These assumptions are consistent with host–parasite systems with strongly seasonal or pulsed transmission, including salmonid diseases and the plant pathosystems discussed above; the full empirical justification is provided in Supplementary Information S4.1.

A susceptible host of genotype *i* produces offspring according to

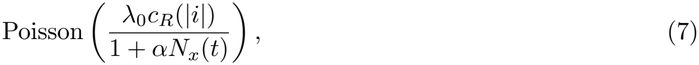

whereas an infected host of genotype *i* carrying parasite genotype *j* produces offspring according to

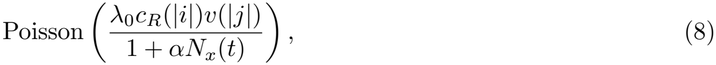

Where *λ* _0_ is baseline fecundity,*α* is the competition coefficient, and *N* _*x*_(*t*) is total host abundance in patch *x* before reproduction. Drawing offspring numbers form a Poisson distribution allows us to incorporate demographic stochasticity (Deshpande et al., 2025a). Parents are removed after reproduction, and newborns inherit their parent’s genotype, with each locus mutating independently with probability *m*_*H*_ in hosts and *m* _*P*_ in parasites.

Newborn hosts are susceptible and are exposed to the local parasite pool. Following a Poisson encounter process (Bailey, 1931; Hassell, 2000; Gustafsson and Sternad, 2007), the probability that a newborn host of genotype*i*in patch*x*escapes infection by all parasite genotypes is

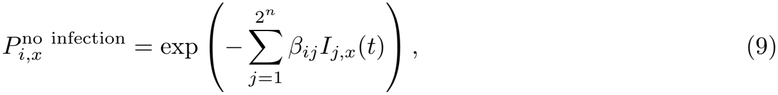

where *I* _*j,x*_(*t*) is the number of infected hosts carrying parasite genotype *j* in patch *x*. Thus, the probability of infection is 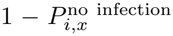. Following Ashby et al. (2014), when infection occurs, the establishing parasite genotype is drawn from the local pool with a probability proportional to its marginal infection probability. This formulation saturates at high number of infected individuals locally and gives rare parasite genotypes an advantage; a comparison with the proportional force-of-infection formulation is provided in Supplementary Information S4.3.

### 4.3 Networks

We compared river-like Optimal Channel Networks (OCNs; Rinaldo et al. 2014), terrestrial-like Random Geometric Graphs (RGGs; Gilarranz and Bascompte 2012), and homogeneous controls with comparable average degree. Because OCNs and RGGs differ in mean degree, we used a regular grid and a ring as homogeneous counterparts. We also generated synthetic networks spanning gradients of degree heterogeneity and assortativity, allowing us to separate the effects of unequal patch degrees from the arrangement of connections among patches. Full network-generation procedures, degree distributions, and network statistics are provided in Supplementary Information S4.4.

#### 4.4 Network Metrics

Degree heterogeneity was quantified using the normalised Herfindahl-Hirschman Index (HHI), which ranges from 0 for regular graphs to 1 for maximally concentrated degree distributions. Assortativity was quantified as Spearman’s rank correlation, *ρ*, between the degrees of connected patches. Formal definitions are provided in Supplementary Information S4.5.1 and S4.5.2.

### 4.5 Simulation protocol

The individual based model was coded in Rust (version 1.97.1). The model was compiled using Cargo (version 1.97.1).

Each patch was initialised with 1,000 hosts, corresponding to one third of the equilibrium density of an uninfected single-patch. Half of the initial hosts were infected, and initial host and parasite genotypes were drawn at random. We ran 20 replicate simulations on independent stochastic network realisations. Case-study simulations ran for 50,000 generations for the networks (Fig. 2, S1, 3, Fig. 1 C-H), and heatmap simulations for a single-patch ran for 100,000 generations (Fig. 1 A,B); the final 200 generations were used for all analysis. Both fixed and varying parameter values are provides in Table S1.

## Author contributions

SN, JD, OK and EAF designed the research. SN setup the code in collaboration with IB. SN analysed the model. SN and EAF interpreted the results. SN, JD and EAF wrote the manuscript. All authors commented on the text.

## Acknowledgments

This work was funded by a grant from the PEPR Prezode via the Agence Nationale de Recherche (ZOOCAM, grant no. ANR-23-PEPZ-0004). This work was carried out within the framework of the Camargue Health-Environment “Zone Atelier” (ZACAM) of the ‘LONG-TERM SOCIO-ECOLOGICAL RESEARCH - LTSER” network (RZA), funded by the French National Center for Scientific Research (CNRS-EE). E.A.F has received financial support from the CNRS through the MITI interdisciplinary program. O.K. was funded by a grant from the Agence Nationale de Recherche (grant no. ANR-20-CE02-0023-01). This is publication ISEM-XXXX-XXX of the Institut des Sciences de l’Evolution - Montpellier.

## AI statement

Deepseek (version 3) was used for assistance with coding and text editing.

## Code and Data availability

Simulation and analysis code is available at XXX.

## Supplementary Information

### S1 Supplementary Theory

#### S1.1 Degree heterogeneity in networks select for reduced parasite infectivity when hosts are resistant

We examine impact of spatial network structure when host resistance is evolves to its maximum and can therefore, no longer evolve to be higher(Fig. S1; white star in Fig. 1). This scenario corresponds to the red region of Fig. 1A, where resistance has reached its upper bound. As expected, host resistance remains un-changed across network types (Fig. S1A). In contrast, parasite infectivity in OCNs exhibits a pronounced U-shaped relationship with dispersal, a pattern absent from the other network types (Fig. S1B).

As in the the case study (Section. 2), this pattern arises from degree-dependent redistribution of hosts across the network (Fig. S12). The higher transmission and virulence costs in this scenario result in lower overall population densities. At low and intermediate dispersal, host abundance increases across the network, whereas at high dispersal, net outflow from single-neighbour patches sharply reduces their density and lowers network-wide abundance (Fig. S1C).

The density-dependent transmission mechanism identified in Section 2.2 also operates here. Highly connected patches accumulate hosts, including infected hosts, through net influx. As infected-host density increases, transmission saturates and the marginal benefit of additional infectivity declines. Consequently, lower-infectivity genotypes gain a relative advantage in highly connected patches, producing a spatial mosaic in which single-degree patches harbour the most infective parasites and highly connected patches the least (Fig. S1D).

We further tested this mechanism in single-patch simulations by increasing infected-host density through higher host intrinsic growth rates (*λ*_0_). As density increased, mean parasite infectivity declined and the relative advantage of lower-infectivity genotypes increased (Fig. S1E; see also Fig. 2H). However, OCN hubs maintain higher infectivity and lower *w* _3_*/w*_4_ ratios than predicted by the single-patch relationship at comparable infected-host densities. This discrepancy likely reflects dispersal-mediated gene flow between patches: hubs export lower-infectivity alleles to low-degree patches, where selection against them is stronger, while importing higher-infectivity alleles from these patches (see flux of 3-infectivity allele parasites in Fig. S30). This coupling constrains the evolutionary response across the metapopulation, although the overall relationship between infected-host density and infectivity remains negative.

RGGs show a weaker version of this mechanism because their degree distributions are less heterogeneous (Fig. S16B; Supplementary Information S29). Their degree-6 patches do not reach sufficiently high infected-host densities to generate strong selection against high infectivity. Accordingly, *w* _3_*/w*_4_ remains close to zero across dispersal rates (Fig. S29D,E).

**Figure S1:**
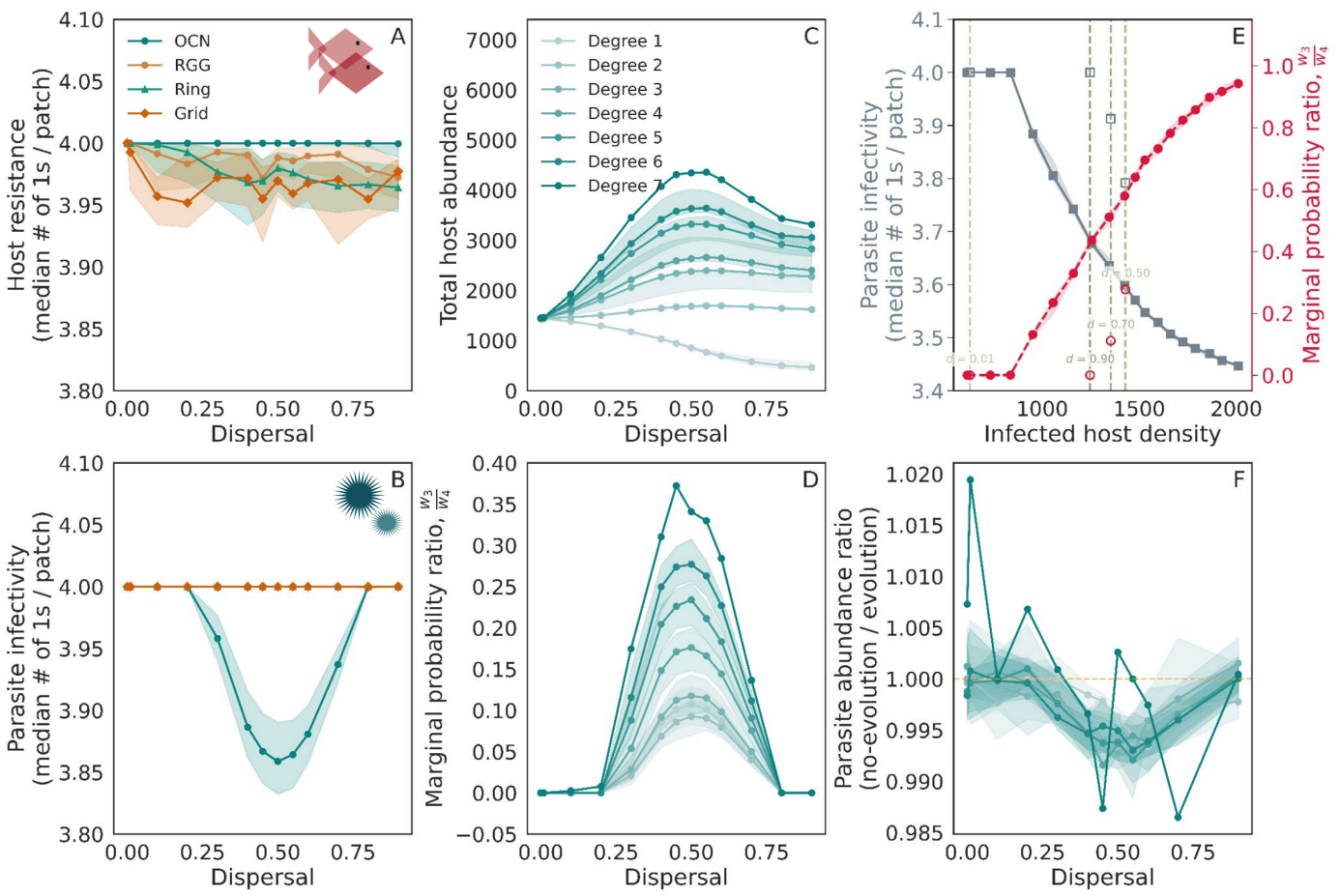
Impact of dispersal and network-type for intermediate virulence and transmission costs (*µ*_*V*_ = 0.5,*µ* _*T*_ = 0.6), with a low resistance cost (*µ*_*R*_ = 0.000256) as shown by the white star in Fig. 1. A) Host resistance (median number of “1” alleles in the host population) per patch as a function of dispersal. Results are shown for four network-types: OCN (teal), RGG (orange), ring (light green), and grid (dark green). Shaded regions indicate the interquartile range (IQR) across patches and replicates. B) Parasite infectivity (median number of “1” alleles in the parasite per patch) as a function of dispersal. C) Median host abundance per patch, grouped by patch degree in the OCN. D) Median ratio of marginal probability of parasite genotypes with 3 ‘1” or infectivity alleles (*w*_3_) to marginal probability of parasite genotypes with 4 “1” alleles (*w*_4_) as defined by Eqn. S7 (also see *w* _*k*_ plotted as functions of local infected densities in Fig. S10, computed from the pre-dispersal state and grouped by patch degree in the OCN. E) Parasite infectivity as a function of equilibrium infected host densities in a single-patch. The solid grey shows the median number of “1” infectivity alleles across replicates, and the shaded band gives the IQR. The solid red line shows the median ratio of marginal probability of parasite genotypes with 3 “1” or infectivity alleles (*w*_3_) to marginal probability of parasite genotypes with 4 ‘1” alleles (*w*_4_) as defined by Eqn. S7 in the single-patch. Vertical dashed lines mark the median equilibrium infected host densities in the OCN degree-6 patches found in the simulations at dispersal rate, *d*= 0.01,0.5,0.7,0.9 computed from the pre-dispersal state. On each vertical line, the square marker is the median infectivity in those patches (left axis) and the circle marker is the median 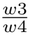 ratio (right axis). F) Ratio of parasite abundance in simulations without evolution, in which host resistance and parasite infectivity are fixed at 1111, to parasite abundance in the full evolutionary simulations for the OCN, grouped by patch degree. Refer to Table S1 for fixed parameters.

To assess whether reduced infectivity is adaptive, we compared parasite abundance in the full evolutionary simulations with abundance in simulations where host resistance and parasite infectivity were fixed at 1111 (Fig. S1F). Parasite abundance was higher in the evolutionary simulations, suggesting that evolved infectivity improves parasite performance relative to a non-evolving maximum-infectivity base-line. However, because host populations are heterogeneous, this comparison provides an indirect measure of adaptation rather than a direct test of *R* _0_ maximisation (Lion and Metz, 2018).

More broadly, our results complement Sato et al. (2024), who showed that metapopulation heterogeneity in host mobility, productivity, and immunocompetence can increase pathogen virulence and infectiousness. We identify an important contrasting outcome: when host resistance is constrained at its upper bound, topology-driven density heterogeneity can instead select for reduced parasite infectivity. This within-network selection mosaic extends prudent-parasite theory to heterogeneous spatial networks, where local density and transmission saturation can favour reduced parasite infectivity (Lion and Boots, 2010).

#### S1.2 Effect of culling on equilibrium prevalence

We consider the host–parasite model with random culling. Only single genotype of host and parasite are present. Let*c*be the fraction of hosts removed each generation, and define the surviving fraction

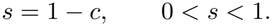

Let *v* be the relative loss of fecundity of the infected therefore (1*−v*).*I* (*t*) gives population contributed by*I* (*t*).

The population dynamic equations are given by

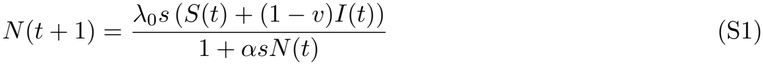

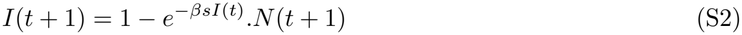

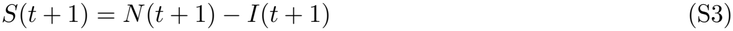

Replacing *S*(*t*) with *N* (*t*)*−I* (*t*) The dynamics of total population size *N* (*t*) and infected number *I*(*t*) are

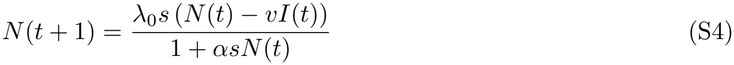

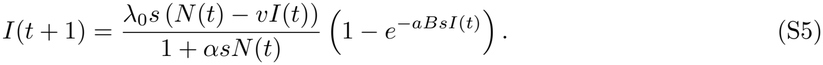

Here *λ* _0_ *>*1 is the intrinsic growth rate,*α* is the competition coefficient, *v* is the virulence (fecundity reduction),*B* is the transmission rate, and *a* is the searching efficiency.

##### Equilibrium equations (fixed point)

At equilibrium *N* (*t*+ 1) =*N* (*t*) =*N* ^*\**^ and *I* (*t*+ 1) =*I* (*t*) =*I* ^*\**^. Define the equilibrium prevalence

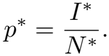

From (1), using *I* ^*\**^ =*p* ^*\**^*N* ^*\**^, we obtain

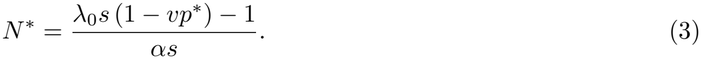

From (2), the equilibrium prevalence satisfies

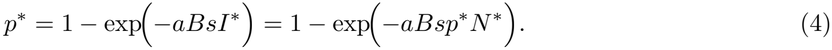

Substituting (3) into (4), let

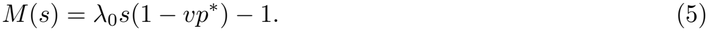

Then *N* ^*\**^ =*M/*(*αs*), and

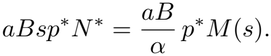

Set

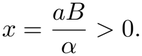

Thus the equilibrium prevalence equation becomes

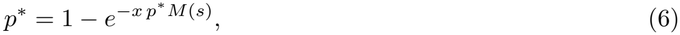

or equivalently,

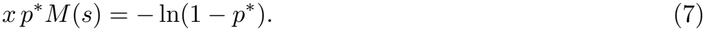

Equation (7) is the key fixed-point relation that determines *p* ^*\**^ as a function of the surviving fraction*s*.

##### Derivative of equilibrium prevalence with respect to *s*

We now differentiate (6) implicitly with respect to *s*. Let *p*=*p* ^*\**^(*s*) for brevity. From (6),

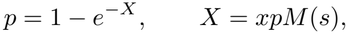

So

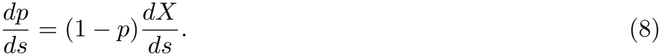

From the definition of *M* in (5),

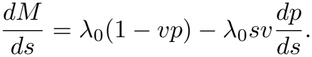

Therefore,

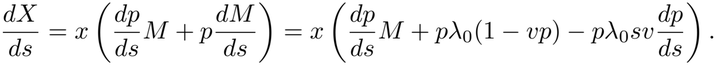

Factor the *dp/ds* term:

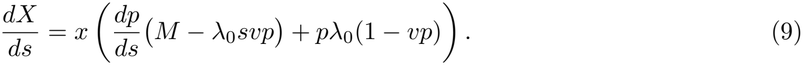

Substitute (9) into (8):

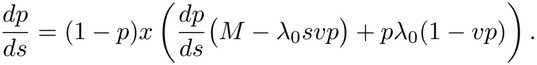

Bring the term containing *dp/ds* to the left-hand side:

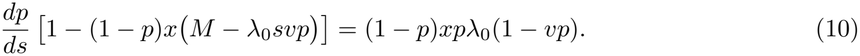

Thus

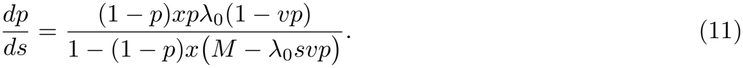

##### Sign of the derivative

We show that the denominator in (11) is strictly positive. From (7),

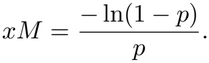

Hence

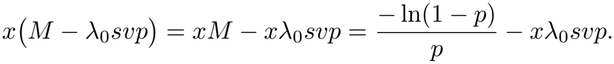

The denominator *D* in (11) becomes

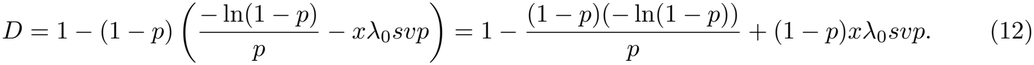

Let

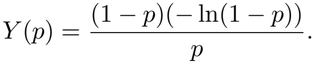

For 0*< p <*1, write *u*=*−*ln (1*−p*)*>*0. Then *p*= 1*−e* ^*−u*^, and

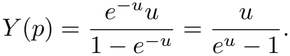

Since*e* ^*u*^ *−*1*> u* for all *u >*0, we have *Y* (*p*)*<*1. Therefore,

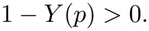

The last term in (12), (1*−p*)*x λ* _0_ *svp*, is strictly positive for *p >*0 (provided *v >*0 and *s >*0). Hence

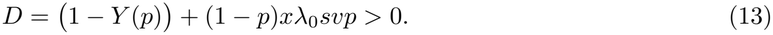

The numerator in (11) is also positive because 0*< p <*1, 1*−vp >*0 (otherwise the population cannot sustain positive growth), and all other parameters are positive. Thus

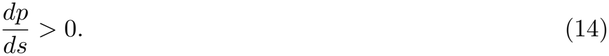

##### Effect of culling fraction *c*

Since *s*= 1*−c*,

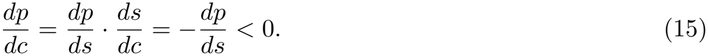

##### Conclusion

Therefore, the equilibrium prevalence *p* ^*\**^ is a strictly decreasing function of the culling fraction *c*:

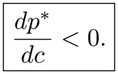

Culling reduces the equilibrium prevalence of the infection.

### S2 Supplementary Tables

**Table S1:**
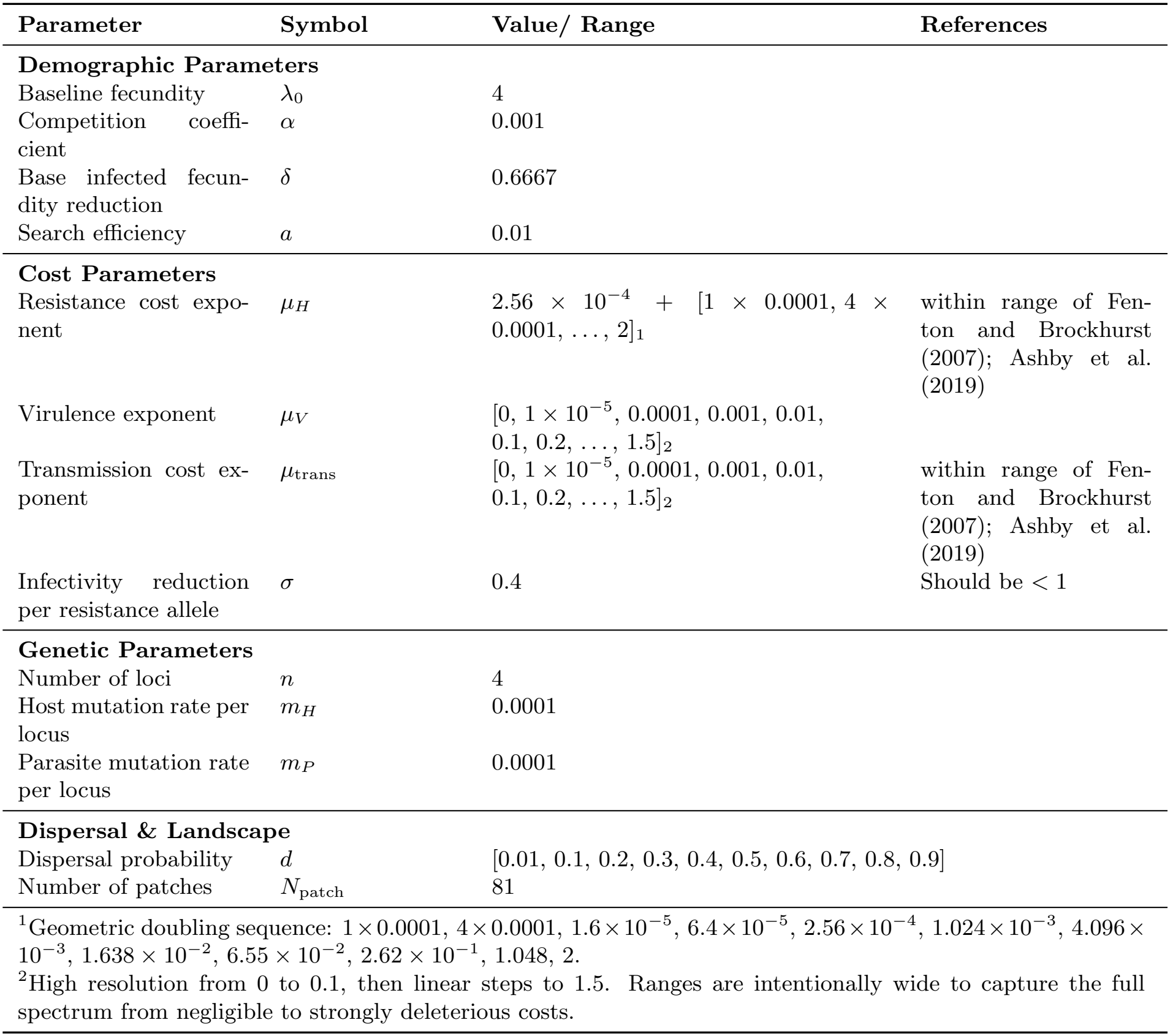
Model parameters and explored ranges.

| Parameter | Symbol | Value/ Range | References |
|---|---|---|---|
| **Demographic Parameters** |  |  |  |
| Baseline fecundity | λ<sub>0</sub> | 4 |  |
| Competition coeffi-cient | α | 0.001 |  |
| Base infected fecun-dity reduction | δ | 0.6667 |  |
| Search efficiency | *a* | 0.01 |  |
| **Cost Parameters** |  |  |  |
| Resistance cost expo-nent | μ<sub>H</sub> | 2.56 × 10<sup>−4</sup> + [1 × 0.0001, 4 × 0.0001, ..., 2]<sub>1</sub> | within range of Fen-ton and Brockhurst (2007); Ashby et al. (2019) |
| Virulence exponent | μ<sub>V</sub> | [0, 1 × 10<sup>−5</sup>, 0.0001, 0.001, 0.01, 0.1, 0.2, ..., 1.5]<sub>2</sub> |  |
| Transmission cost ex-ponent | μ<sub>trans</sub> | [0, 1 × 10<sup>−5</sup>, 0.0001, 0.001, 0.01, 0.1, 0.2, ..., 1.5]<sub>2</sub> | within range of Fen-ton and Brockhurst (2007); Ashby et al. (2019) |
| Infectivity reduction per resistance allele | σ | 0.4 | Should be < 1 |
| **Genetic Parameters** |  |  |  |
| Number of loci | *n* | 4 |  |
| Host mutation rate per locus | *m*<sub>H</sub> | 0.0001 |  |
| Parasite mutation rate per locus | *m*<sub>P</sub> | 0.0001 |  |
| **Dispersal & Landscape** |  |  |  |
| Dispersal probability | *d* | [0.01, 0.1, 0.2, 0.3, 0.4, 0.5, 0.6, 0.7, 0.8, 0.9] |  |
| Number of patches | *N*<sub>patch</sub> | 81 |  |

<sup>1</sup>Geometric doubling sequence: 1 × 0.0001, 4 × 0.0001, 1.6 × 10<sup>−5</sup>, 6.4 × 10<sup>−5</sup>, 2.56 × 10<sup>−4</sup>, 1.024 × 10<sup>−3</sup>, 4.096 × 10<sup>−3</sup>, 1.638 × 10<sup>−2</sup>, 6.55 × 10<sup>−2</sup>, 2.62 × 10<sup>−1</sup>, 1.048, 2.
<sup>2</sup>High resolution from 0 to 0.1, then linear steps to 1.5. Ranges are intentionally wide to capture the full spectrum from negligible to strongly deleterious costs.

### S3 Supplementary Figures

**Figure S2:**
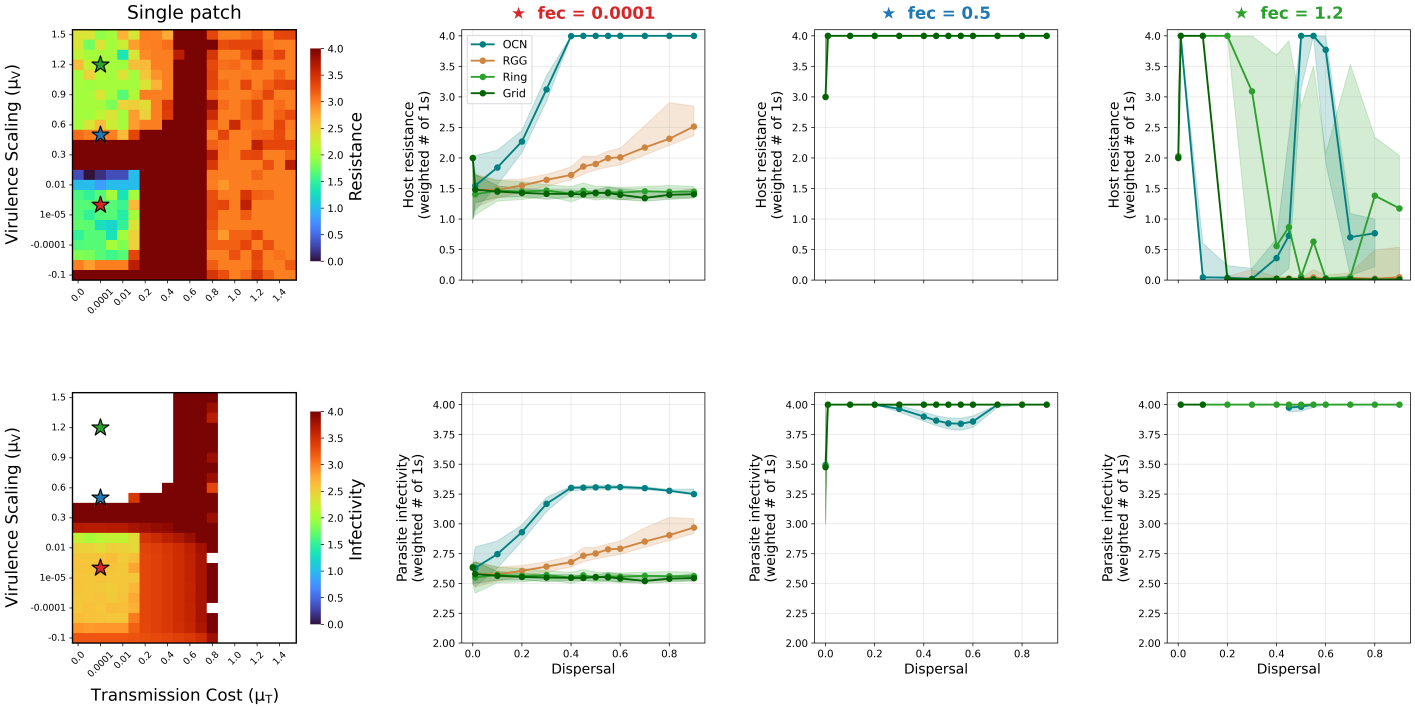
Host-parasite coevolution at low transmission cost scaling*µ* _*T*_ = 0.0001 with resistance cost scaling*µ* _*R*_ = 0.000256 and dispersal rate*d*= 0.5. The leftmost column shows single-patch heatmaps of evolved host resistance (top; median number of “1” alleles per patch) and parasite infectivity (bottom; median number of “1” alleles per patch) across the parameter space of transmission cost scaling (*µ*_*T*_ ; x-axis) and virulence scaling (*µ*_*V*_ ; y-axis). Colour intensity ranges from blue (low) to yellow (high) on a fixed 0–4 scale. White cells indicate the absence of the parasite. The three coloured stars mark the specific fecundity cost values used in the neighbouring columns. Columns 2–4 show the median number of “1” alleles in the host (top row) and parasite (bottom row) per patch as a function of dispersal, for the three fecundity costs highlighted by the stars (red, blue, green corresponding to*µ* _*F*_ = 0.0001,0.5,1.2 respectively). Results for the OCNs are shown in teal, the RGG in orange, the ring network in vermilion and the grid network in sea green; all networks have 81 patches and the shaded region gives the IQR across patches. Refer to Table S1 for fixed parameters.

**Figure S3:**
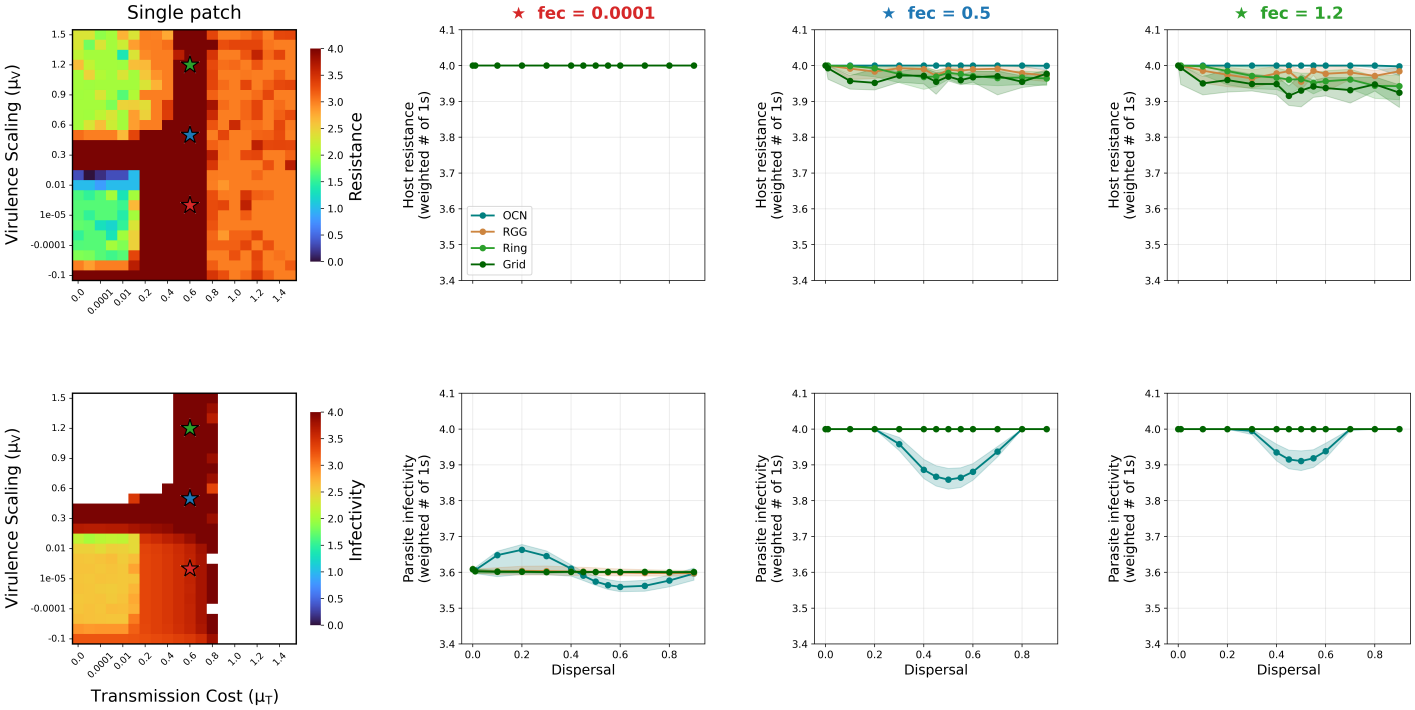
Host-parasite coevolution at low transmission cost scaling*µ* _*T*_ = 0.6 with resistance cost scaling*µ* _*R*_ = 0.000256 and dispersal rate*d*= 0.5. The leftmost column shows single-patch heatmaps of evolved host resistance (top; median number of “1” alleles per patch) and parasite infectivity (bottom; median number of “1” alleles per patch) across the parameter space of transmission cost scaling (*µ*_*T*_ ; x-axis) and virulence scaling (*µ*_*V*_ ; y-axis). Colour intensity ranges from blue (low) to yellow (high) on a fixed 0–4 scale. White cells indicate the absence of the parasite. The three coloured stars mark the specific fecundity cost values used in the neighbouring columns. Columns 2-4 show the median number of “1” alleles in the host (top row) and parasite (bottom row) per patch as a function of dispersal, for the three fecundity costs highlighted by the stars (red, blue, green corresponding to*µ* _*F*_ = 0.0001,0.5,1.2 respectively). Results for the OCNs are shown in teal, the RGG in orange, the ring network in vermilion and the grid network in sea green; all networks have 81 patches and the shaded region gives the IQR across patches. Refer to Table S1 for fixed parameters.

**Figure S4:**
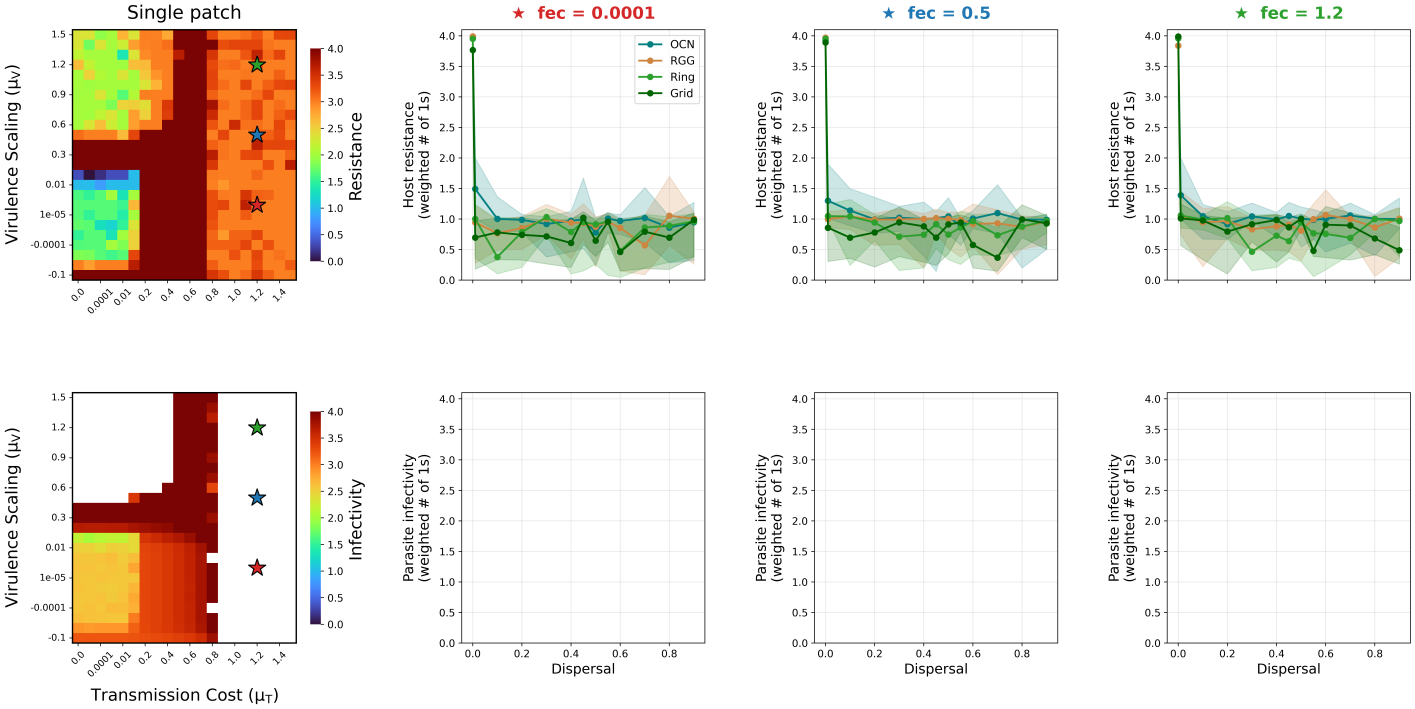
Host-parasite coevolution at low transmission cost scaling*µ* _*T*_ = 1.2 with resistance cost scaling*µ* _*R*_ = 0.000256 and dispersal rate*d*= 0.5. The leftmost column shows single-patch heatmaps of evolved host resistance (top; median number of “1” alleles per patch) and parasite infectivity (bottom; median number of “1” alleles per patch) across the parameter space of transmission cost scaling (*µ*_*T*_ ; x-axis) and virulence scaling (*µ*_*V*_ ; y-axis). Colour intensity ranges from blue (low) to yellow (high) on a fixed 0–4 scale. White cells indicate the absence of the parasite. The three coloured stars mark the specific fecundity cost values used in the neighbouring columns. Columns 2-4 show the median number of “1” alleles in the host (top row) and parasite (bottom row) per patch as a function of dispersal, for the three fecundity costs highlighted by the stars (red, blue, green corresponding to*µ* _*F*_ = 0.0001,0.5,1.2 respectively). Results for the OCNs are shown in teal, the RGG in orange, the ring network in vermilion and the grid network in sea green; all networks have 81 patches and the shaded region gives the IQR across patches. Refer to Table S1 for fixed parameters.

**Figure S5:**
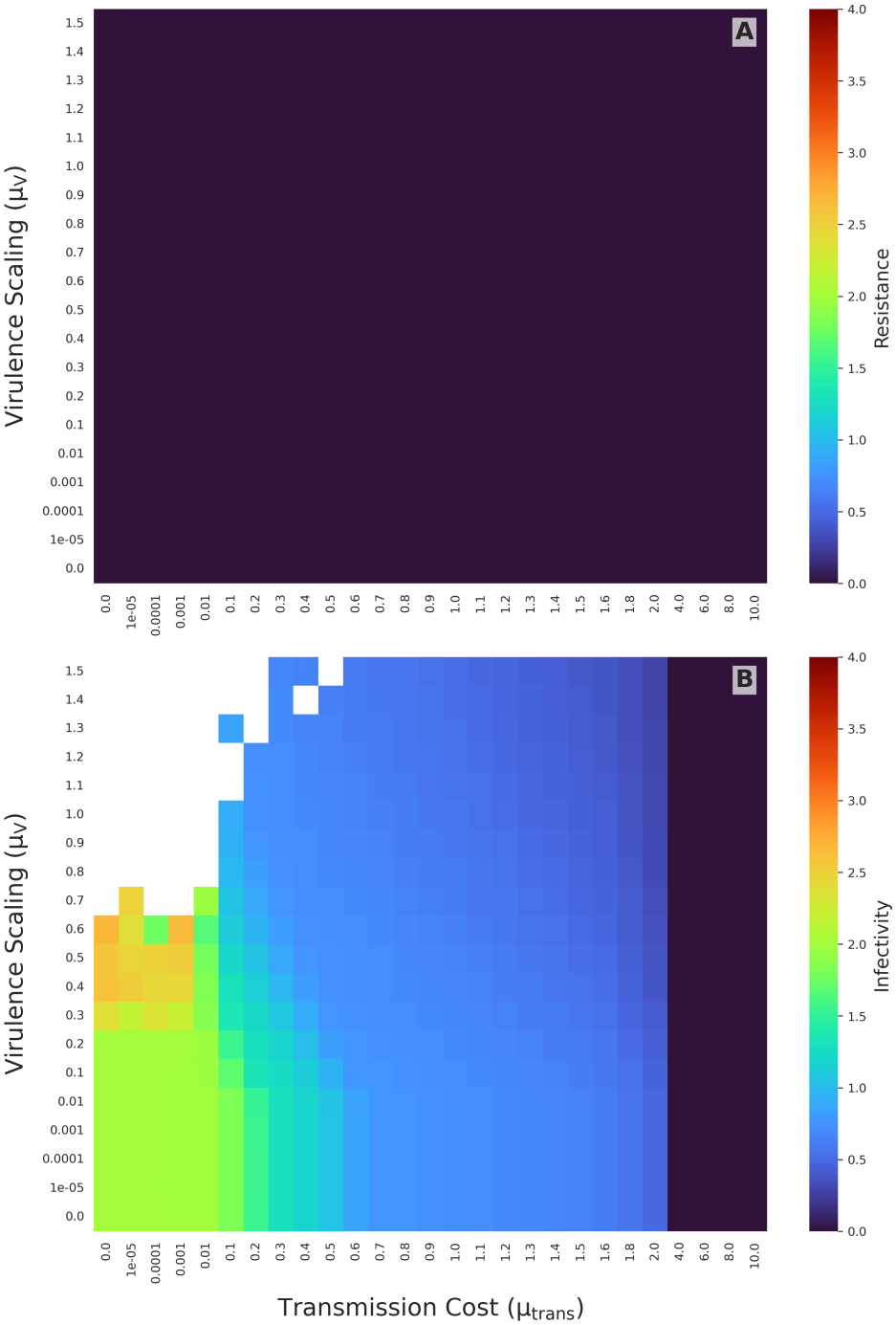
Phase diagram exploring the effects of transmission cost scaling (*µ*_*trans*_; y-axis) and virulence scaling (*µ*_*V*_ ; x-axis) on host–parasite coevolution in a single patch when resistance cost*µ* _*H*_ = 0.25. The top graph A) shows evolved host resistance, quantified as the median number of resistance alleles (‘1’s) across the host population, while the bottom graph B) shows evolved parasite infectivity, quantified as the median number of infectivity alleles (‘1’s) per patch. Colour intensity ranges from blue (low) to red (high), with warmer colours indicating greater evolutionary investment in resistance or infectivity.

**Figure S6:**
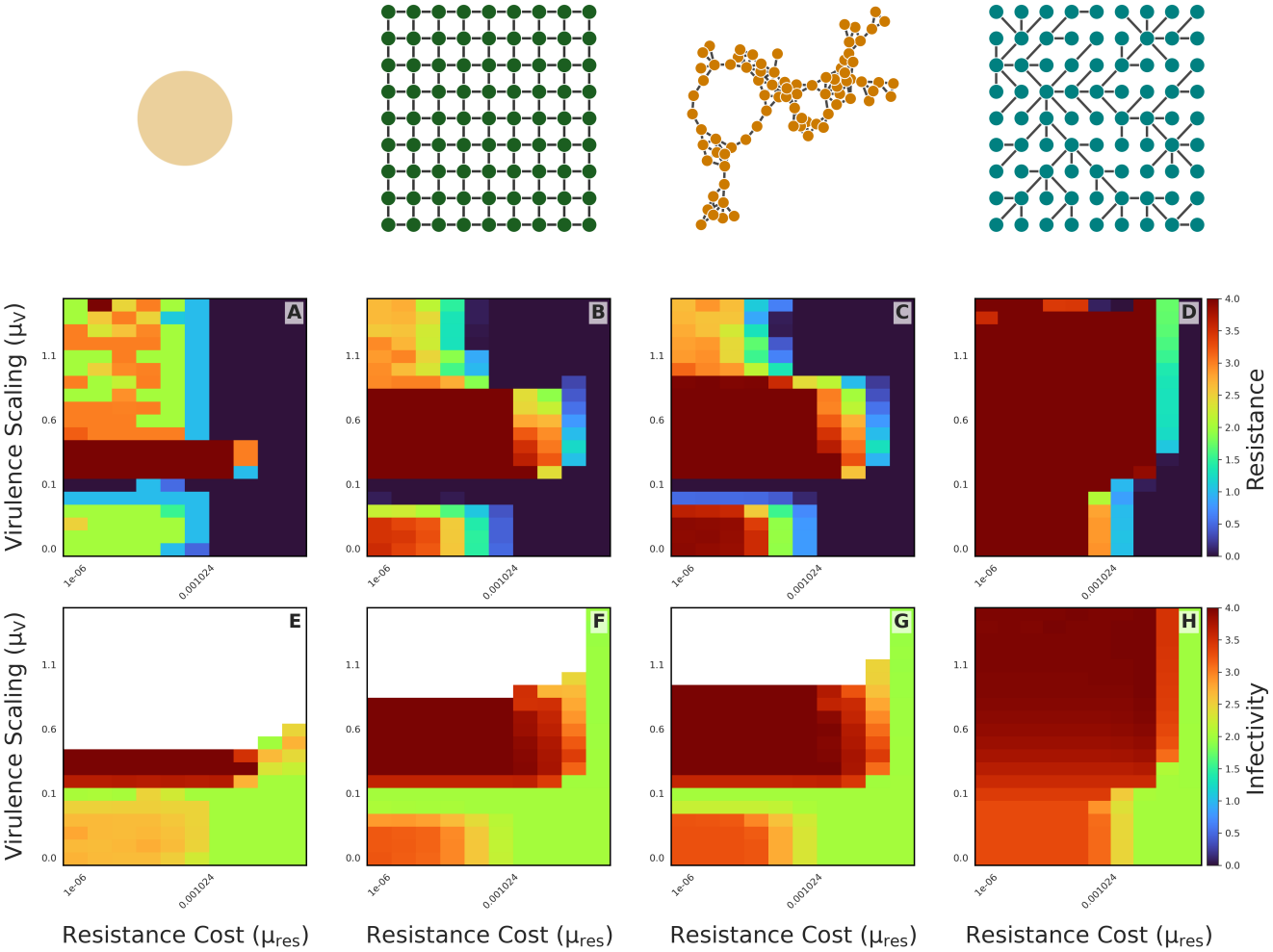
Phase diagram exploring the effects of virulence scaling (*µ*_*V*_ ; y-axis) and resistance cost scaling (*µ*_*H*_ ; x-axis) on host-parasite coevolution across four landscape types when transmission cost scaling *µ*_*trans*_ = 0. Each pair of panels corresponds to a specific spatial structure: (A, B) a single, well-mixed patch; (C, D) a homogeneous grid (torus); (E, F) terrestrial networks modelled as Random Geometric Graphs (RGGs); and (G, H) riverine networks modelled as Optimal Channel Networks (OCNs). The top row (A, C, E, G) shows evolved host resistance, quantified as the median number of resistance alleles (‘1’s) per patch across the host population, while the bottom row (B, D, F, H) shows evolved parasite infectivity, quantified as the median number of infectivity alleles (‘1’s) per patch. Colour intensity ranges from blue (low) to red (high), with warmer colours indicating greater evolutionary investment in resistance or infectivity. Dispersal rate is 0.5.

**Figure S7:**
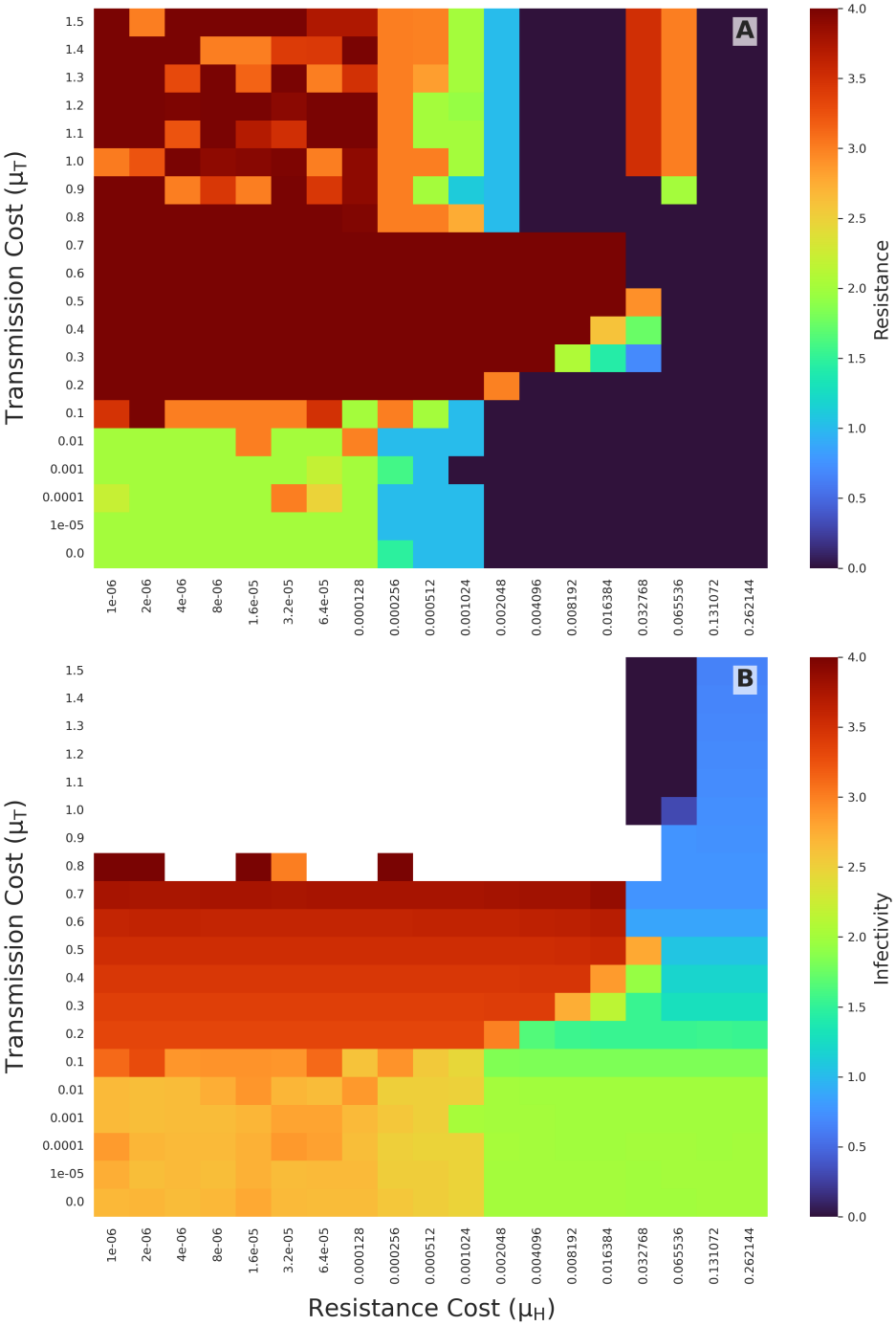
Phase diagram exploring the effects of transmission cost scaling (*µ*_*T*_ ; y-axis) and resistance cost scaling (*µ*_*H*_ ; x-axis) on host-parasite coevolution in a single patch. The top graph A) shows evolved host resistance, quantified as the median number of resistance alleles (‘1’s) across the host population, while the bottom graph B) shows evolved parasite infectivity, quantified as the median number of infectivity alleles (‘1’s) per patch. Colour intensity ranges from blue (low) to red (high), with warmer colours indicating greater evolutionary investment in resistance or infectivity.

**Figure S8:**
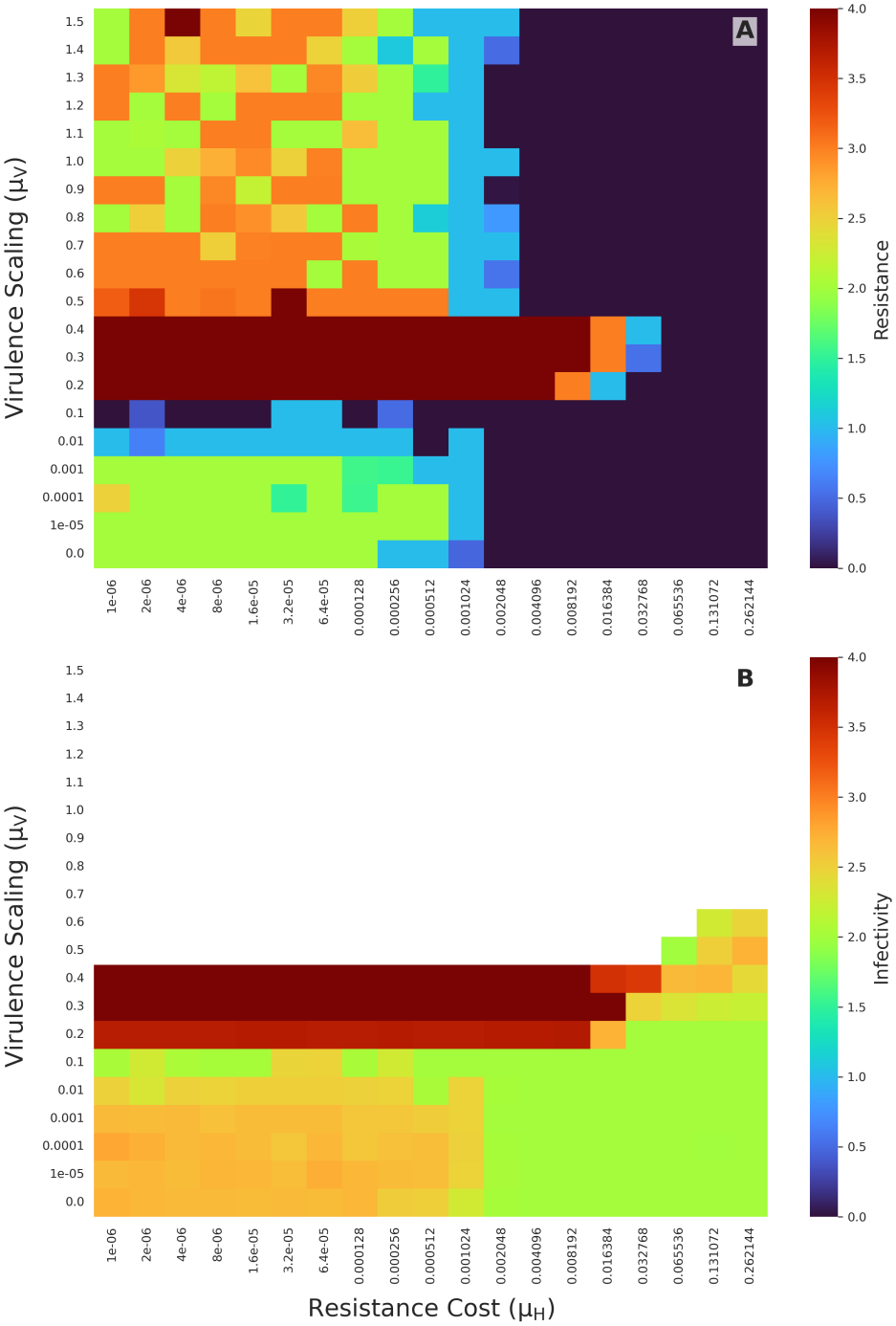
Phase diagram exploring the effects of virulence scaling (*µ*_*V*_ ; y-axis) and resistance cost scaling (*µ*_*H*_ ; x-axis) on host-parasite coevolution in a single patch. The top graph A) shows evolved host resistance, quantified as the median number of resistance alleles (‘1’s) across the host population, while the bottom graph B) shows evolved parasite infectivity, quantified as the median number of infectivity alleles (‘1’s) per patch. Colour intensity ranges from blue (low) to yellow (high), with warmer colours indicating greater evolutionary investment in resistance or infectivity.

**Figure S9:**
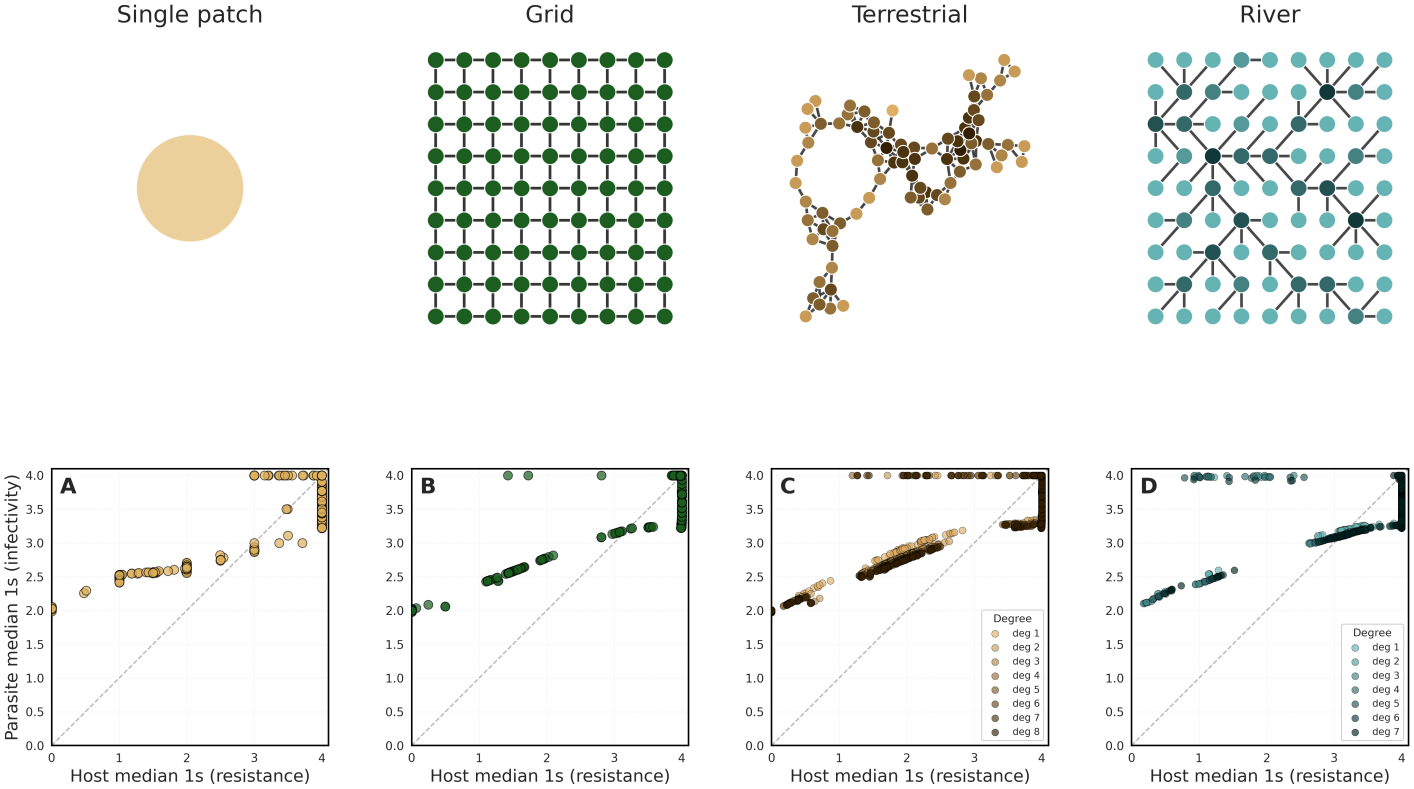
Correlation between coevolved infectivity and resistance alleles for all parameter combinations of transmission cost scaling (*µ*_*T*_ ; x-axis) and virulence scaling (*µ*_*V*_ ; y-axis) as plotted in Fig. 1 for each landscape type. Each panel corresponds to a specific spatial structure: (A) a single, well-mixed patch; (B) a homogeneous grid (torus); (C) terrestrial-like networks modelled as Random Geometric Graphs (RGGs) with results split by patch-degree; and (D) river-like networks modelled as Optimal Channel Networks (OCNs) with results split by patch-degree.

**Figure S10:**
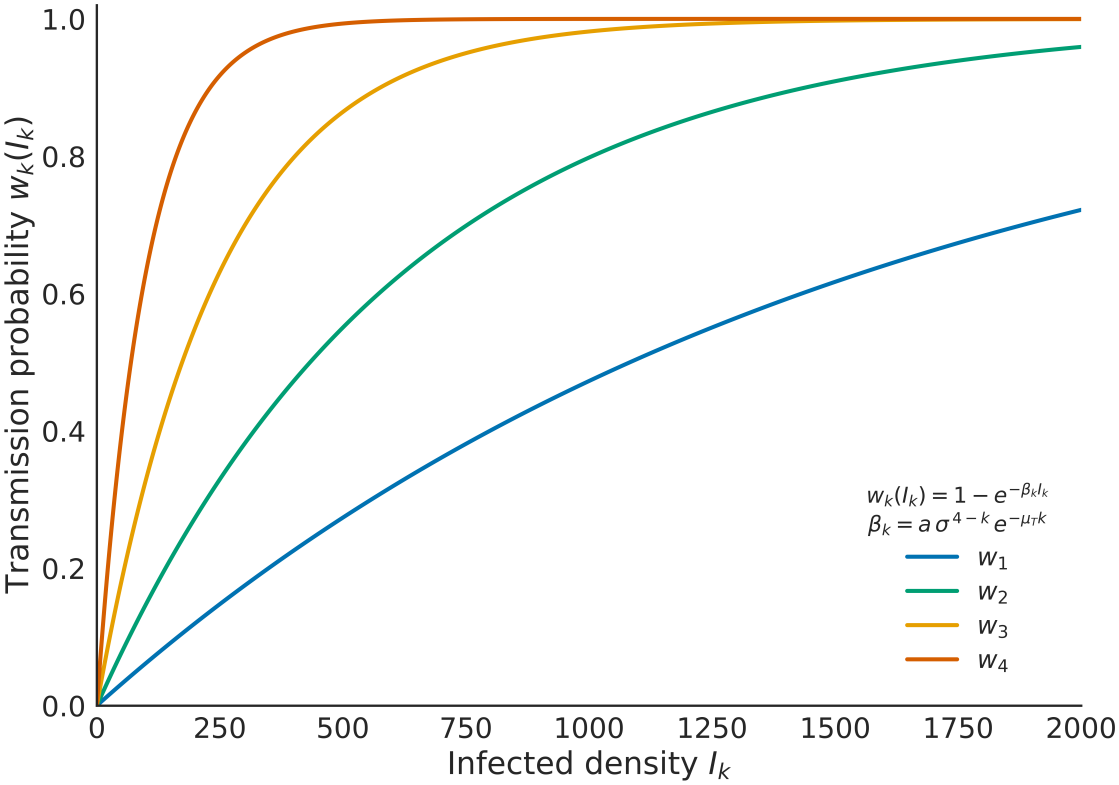
Analytical relationship between the transmission probability 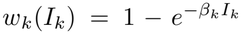 of a parasite genotype carrying *k∈ {*1,2,3,4*}*infectivity alleles and the local density of hosts infected by that genotype, *I* _*k*_. Curves are evaluated from Eqn. 6 for a maximally resistant host, so that the number of effective resistance loci is *d* _*ij*_ = 4*−k*and 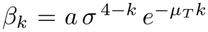. All curves use the default parameters *σ*= 0.4,*µ* _*T*_ = 0.0001 and *a*= 0.01. Because *β* _1_ *< β*_2_ *< β*_3_ *< β*_4_, higher infectivity genotypes saturate at lower infected densities and the ratio *w* _*k*_*/w*_4_ increases with *I* for every *k <*4. This is the mechanism underlying the increase of *w* _2_*/w*_4_ (Fig. 2H) and *w* _3_*/w*_4_ (Fig. S1D,E) with local infected density discussed in the main text, and it drives selection against high-infectivity genotypes in high-density patches. Refer to Table S1 for the full list of fixed simulation parameters.

**Figure S11:**
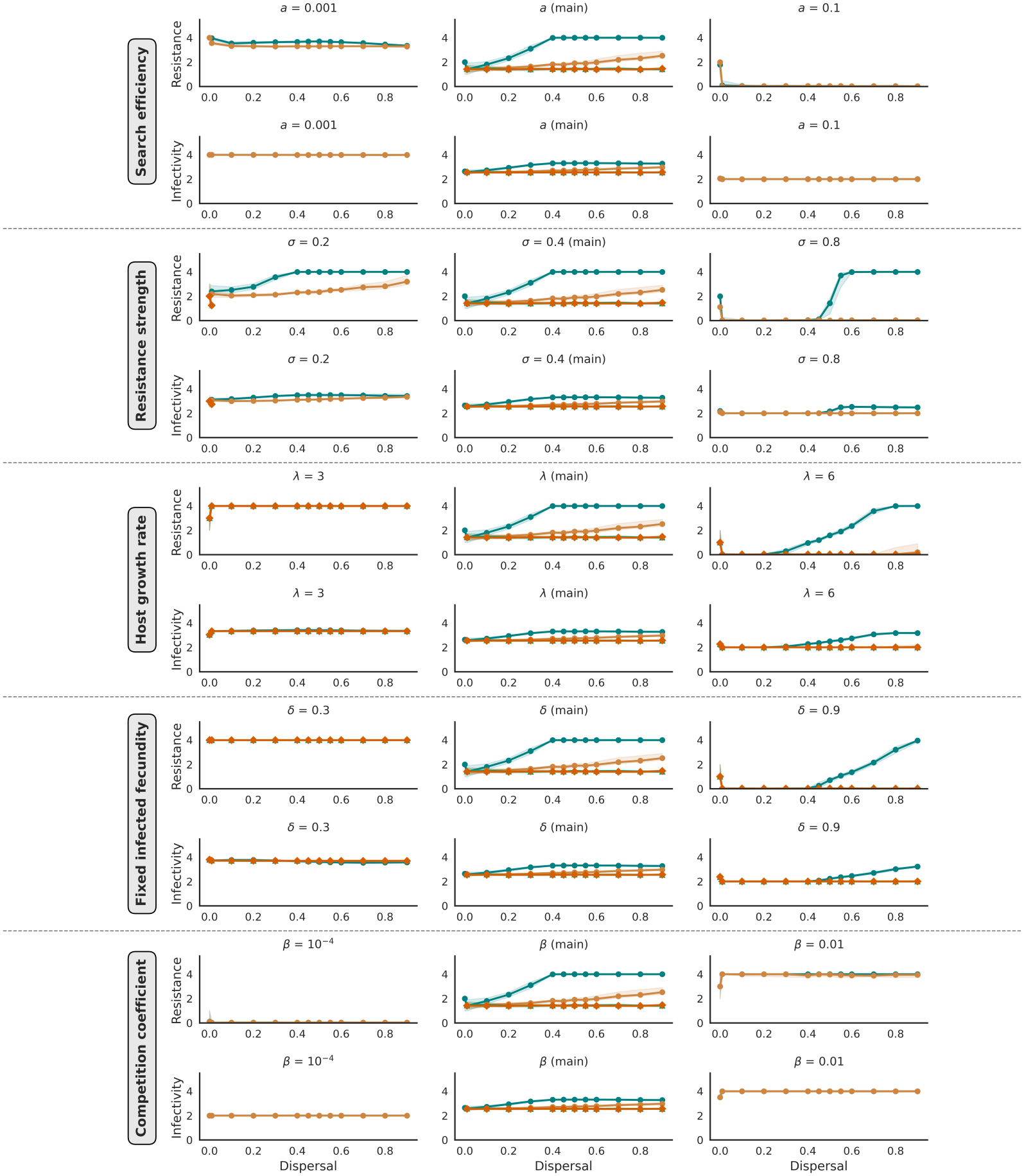
Sensitivity analysis of key ecological and evolutionary parameters on the evolution of host resistance and parasite infectivity across dispersal at fecundity*µ* _*V*_ = 0.0001, transmission cost scaling *µ*_*T*_ = 0.0001, and resistance cost scaling *µ* _*R*_ = 0.000256. Each parameter is represented by a block of two rows: the top row shows the median number of “1” alleles (resistance) in the host population per patch as a function of dispersal, while the bottom row shows the median number of “1” alleles (infectivity) in the parasite per patch. Within each block, the three columns correspond to the lowest, baseline, and highest values of the parameter being varied. From top to bottom, the parameters explored are the search efficiency*a*, the resistance strength*σ*, the fixed infected fecundity *δ*, the host growth rate*λ*, and the competition coefficient *β*. Results for the OCNs are shown in teal, the RGG in orange, the ring network in vermilion, and the grid network in sea green; all networks have 81 patches and the shaded region gives the IQR across patches and replicates. Refer to Table S1 for fixed parameters.

**Figure S12:**
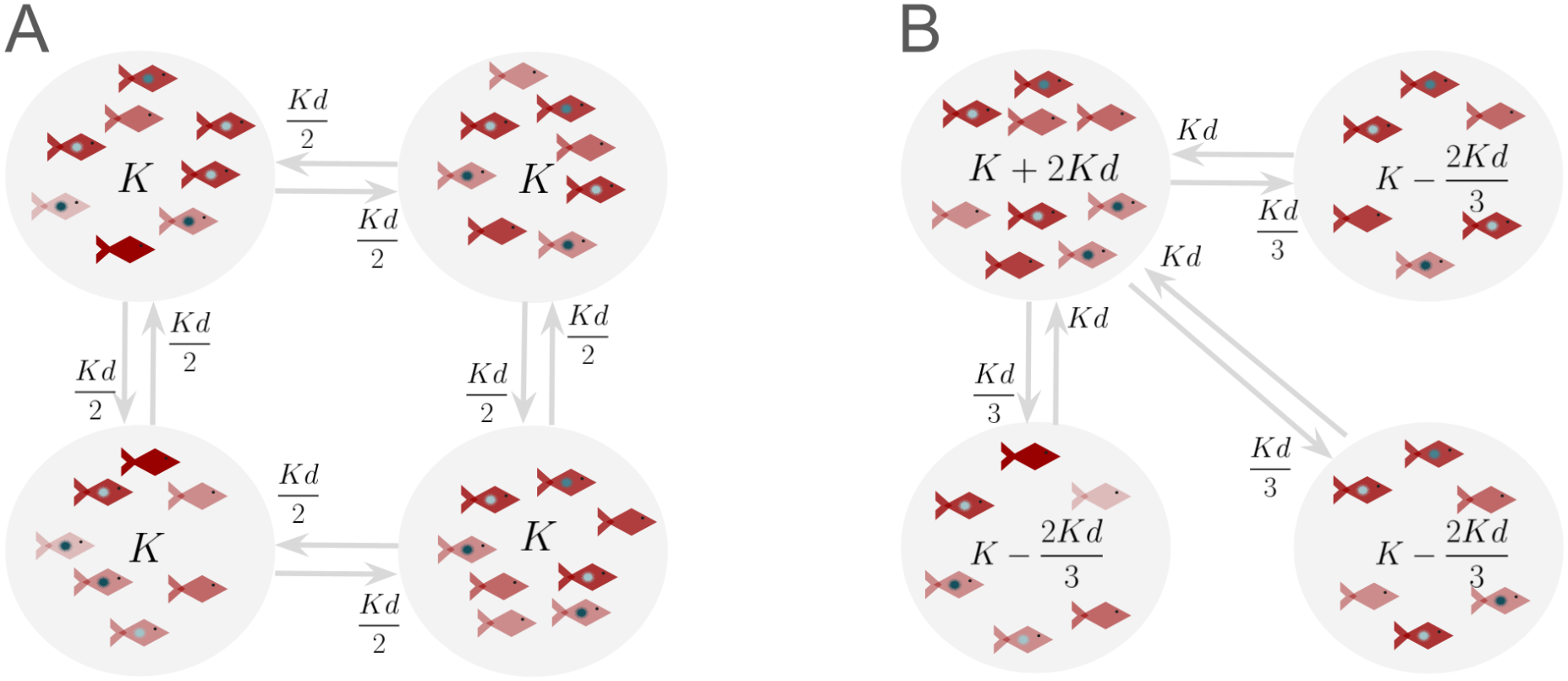
Degree-heterogeneity in a metapopulation leads to density-heterogeneity which is exaggerated in disassortative networks. Our metapopulation assumes in absence of dispersal, each patch maintains the same density, say*K*individuals, and we assume a fixed dispersal rate*d*. Panel A) In a regular graph where all patches have the same degree, number of dispersers entering, in our example it is 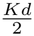 as it is a 2-regular graph, are also the number of dispersers leaving. Hence the density maintained in each patch is *K*. Panel B) In a degree-heterogenous graph, a high degree patch connected to low degree patches receives a net influx of individuals. In our example the net density of the degree-3 patch is *K*(1*−d*) + 3*Kd*=*K*+ 2*Kd*. The degree-1 patches on the other hand, as they are connected to the degree-3 patch, experience a net outflux, in our example it is 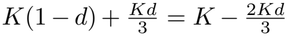. Therefore more disassortative networks, where higher degree patches are more likely to be connected to lower degree patches, maintain greater degree-heterogeneity.

**Figure S13:**
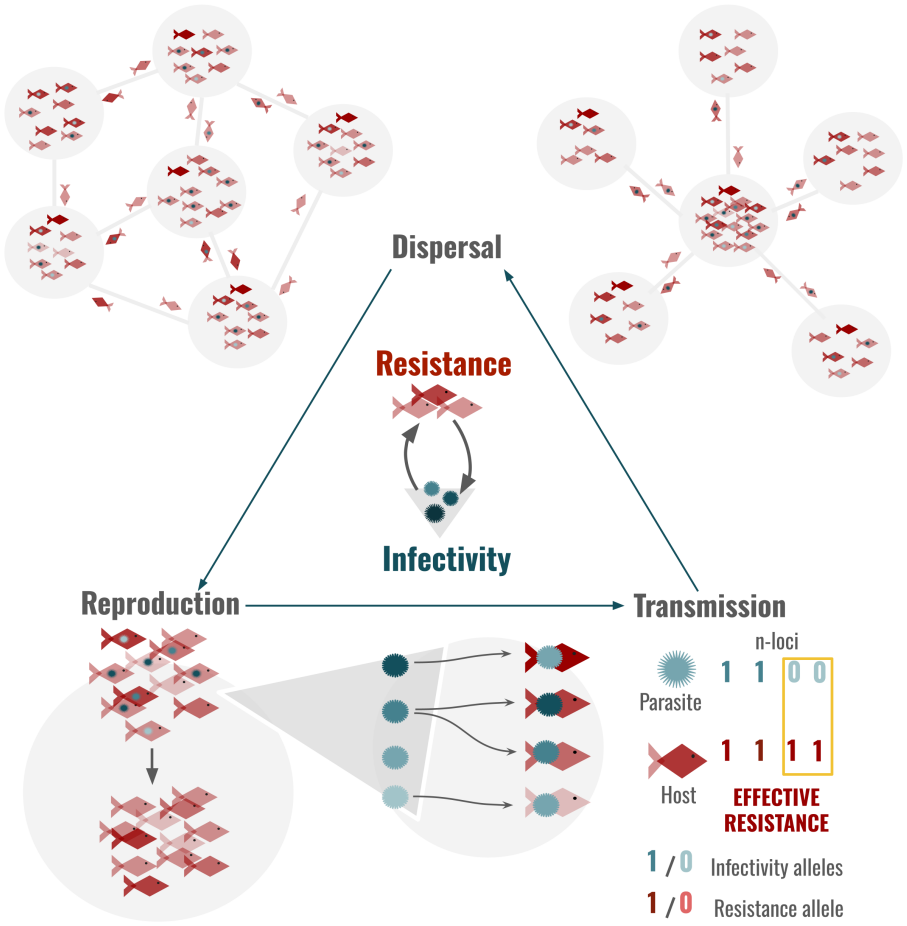
Life cycle of the system has three stages: 1) Dispersal: hosts can disperse with fixed probabilities to their neighbouring patches; 2) Reproduction: Hosts reproduce locally following Beverton-Holt density dependence and number of offsprings are picked from a Poisson distribution for each individual, taking into account cost of resistance and infection status; 3) Transmission: After reproduction, all the offsprings are exposed to pooled local parasites from the parent generation. Effective resistance of a host to a give parasite sequence is determined by pairs of loci in interacting genotypes where the host has a 1 allele and the parasite has a 0 allele (Sasaki, 2000). For example if a parasite sequence of 1100 is attempting to infect a host sequence of 1111 then the 3rd and 4th position of the host will provide it resistance but not its 1st and 2nd position. The 1st position is an effective resistance locus. Each effective resistance locus reduces transmission rate by a factor *σ*. Total probability of encounter with any parasite is first used to determine if infection occurs and if it does, the parasite sequence that would infect is chosen through probability draw weighted by their transmission probabilities on each host.

**Figure S14:**
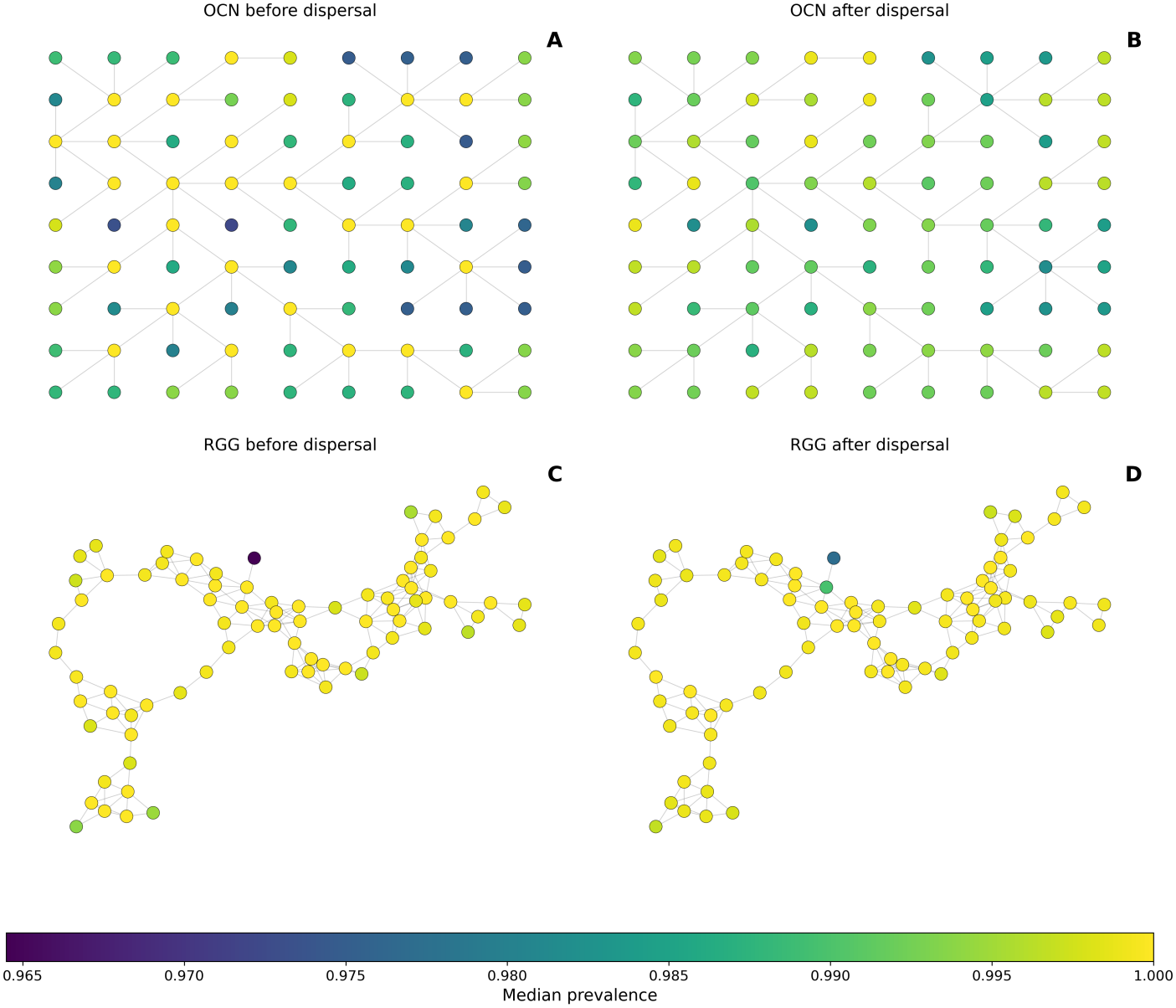
Prevalence distribution in space with *d*= 0.6, when virulence scaling *µ* _*V*_ = 0.0001, transmission cost scaling *µ* _*T*_ = 0.0001 and resistance cost scaling *µ* _*R*_ = 0.000256.The landscape visualise is from a single replicate of the OCN and RGG. Gradient for prevalence is given at the bottom. Panel A) OCN landscape with median prevalence before dispersal; Panel B) OCN landscape with median prevalence after dispersal; Panel C) RGG landscape with median prevalence before dispersal; Panel D) RGG landscape with median prevalence after dispersal. Refer to Table S1 for fixed parameters.

**Figure S15:**
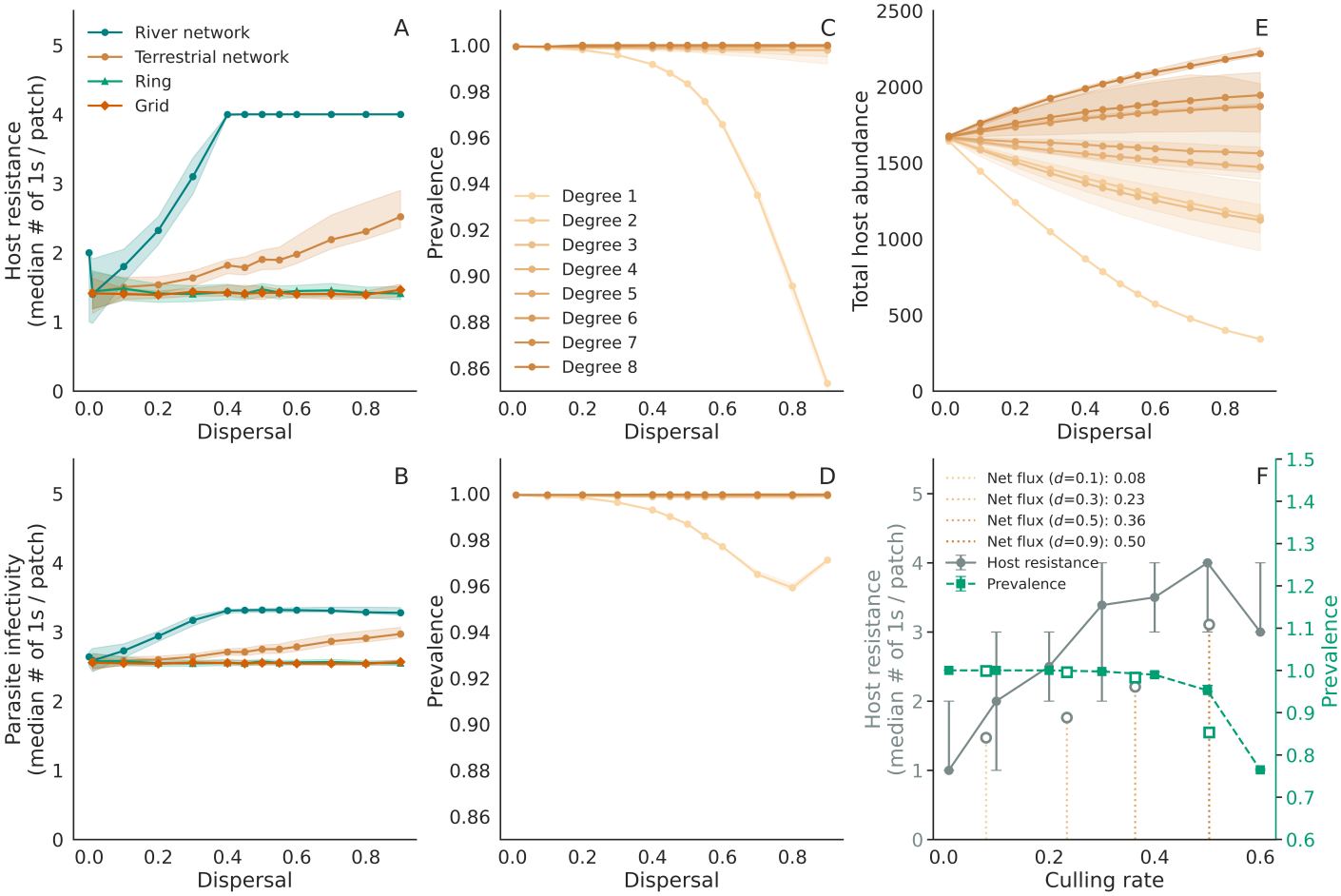
Case study on impact of dispersal and landscape type when virulence scaling *µ* _*V*_ = 0.0001, transmission cost scaling *µ* _*trans*_ = 0.0001 and resistance cost scaling *µ* _*H*_ = 0.000256. Panel A) Median number of 1s in the host per patch with respect to dispersal. Results for the River networks are shown in teal, the Terrestrial networks in orange, the ring and grid network in vermillion and sea green respectively. All networks have 81 patches. Shaded region gives IQR across patches.; B) Median number of 1s in the parasite per patch with respect to dispersal. Shaded region gives IQR across patches.; C) Median prevalence before dispersal per patch grouped by their degrees in the Terrestrial networks. Shaded region gives the IQR across patches. ; D) Median prevalence after dispersal per patch grouped by their degrees in the Terrestrial networks. Shaded region gives the IQR across patches.; E) Median host abundance after dispersal per patch grouped by their degrees in the Terrestrial networks. Shaded region gives the IQR across patches.; F) Simulation results from a single patch with median number of “1” alleles in the host on the y-axis on the left, prevalence in the patch on the right y-axis and increasing culling rate on the x-axis. The vertical lines are drawn at median effective culling experienced through net outflux in the single-neighbour patches of the OCNs at dispersal rates 0.1,0.3,0.5,0.9. The unfilled markers correspond to the median prevalence and resistance experienced the single-neighbour patches. Refer to Table S1 for fixed parameters.

**Figure S16:**
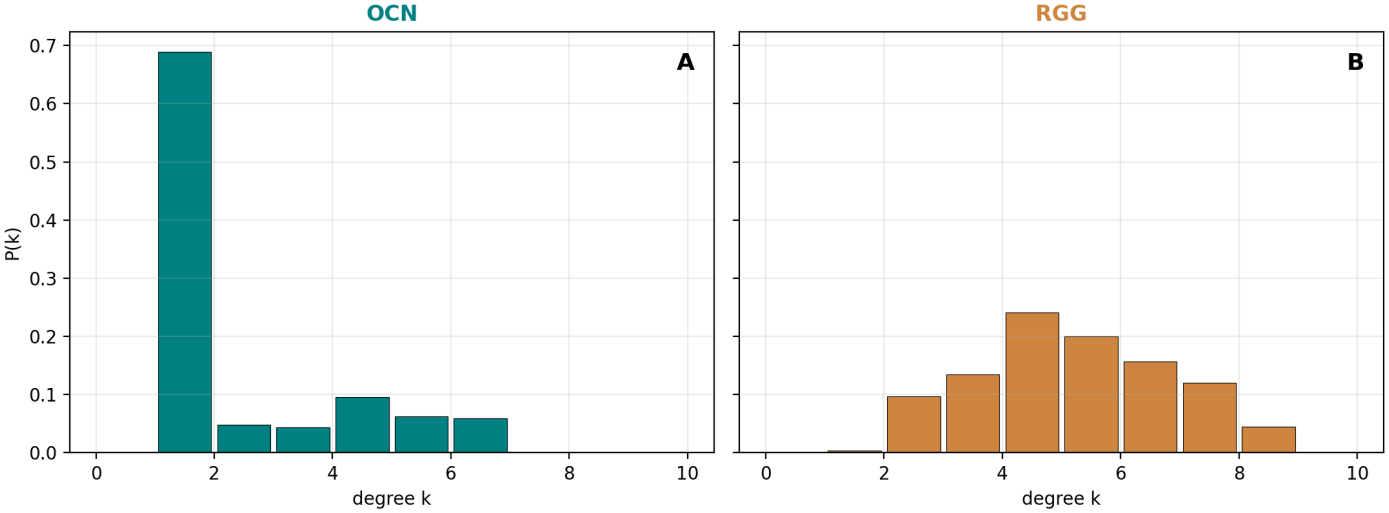
Degree distributions of patches pooled across all 20 replicate networks. A) OCN networks. RGG networks. Bars show the normalized frequency *P* (*k*) of nodes with degree *k*.

**Figure S17:**
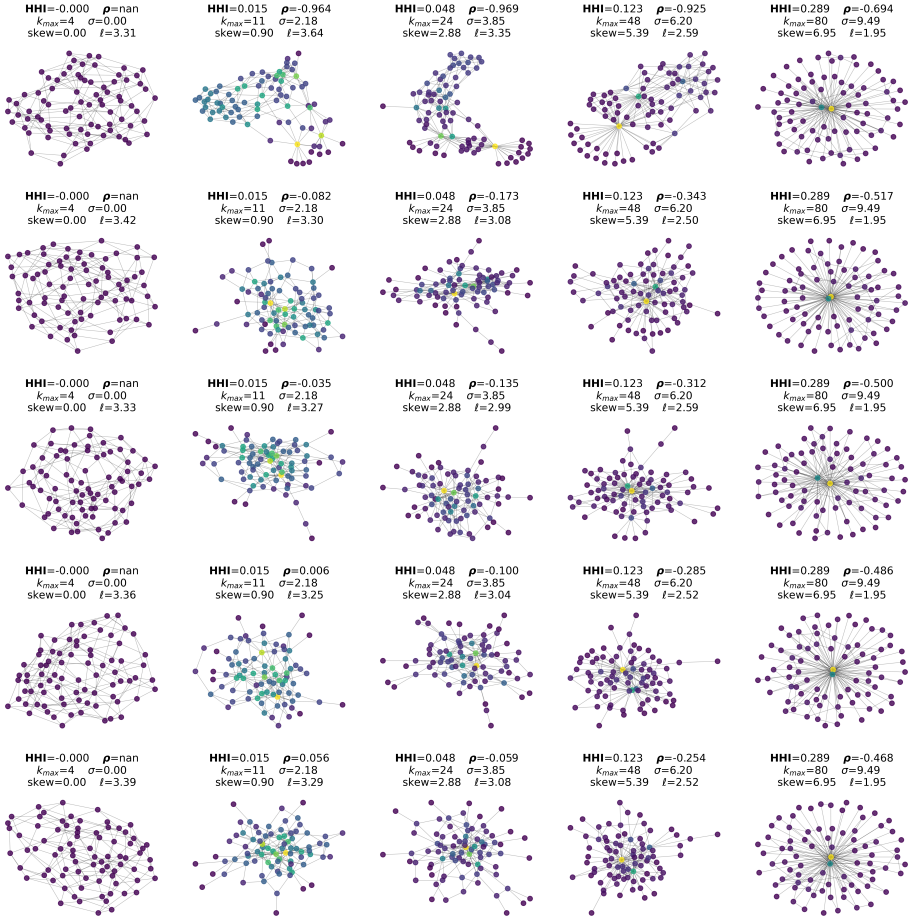
5*×* 5 networks sampled from the 20 *×*20 nested HHI-Spearman’s*ρ*sweep (see algorithm in Supplementary Information S4.6). Columns share the same HHI, which increases from left to right; the Spearman’s *ρ* (assortativity), which increases from top to bottom, while keep HHI constant. First column shows the 4-regular graph with 81 nodes we start the algorithm with hence Spearman’s *ρ* is undefined. Nodes are coloured by degree (dark = low degree, bright = high degree). The label above each panel reports the normalised HHI, Spearman’s *ρ*, maximum degree *k* _max_, degree standard deviation *σ*, degree-distribution skewness, and average shortest-path length *ℓ*.

**Figure S18:**
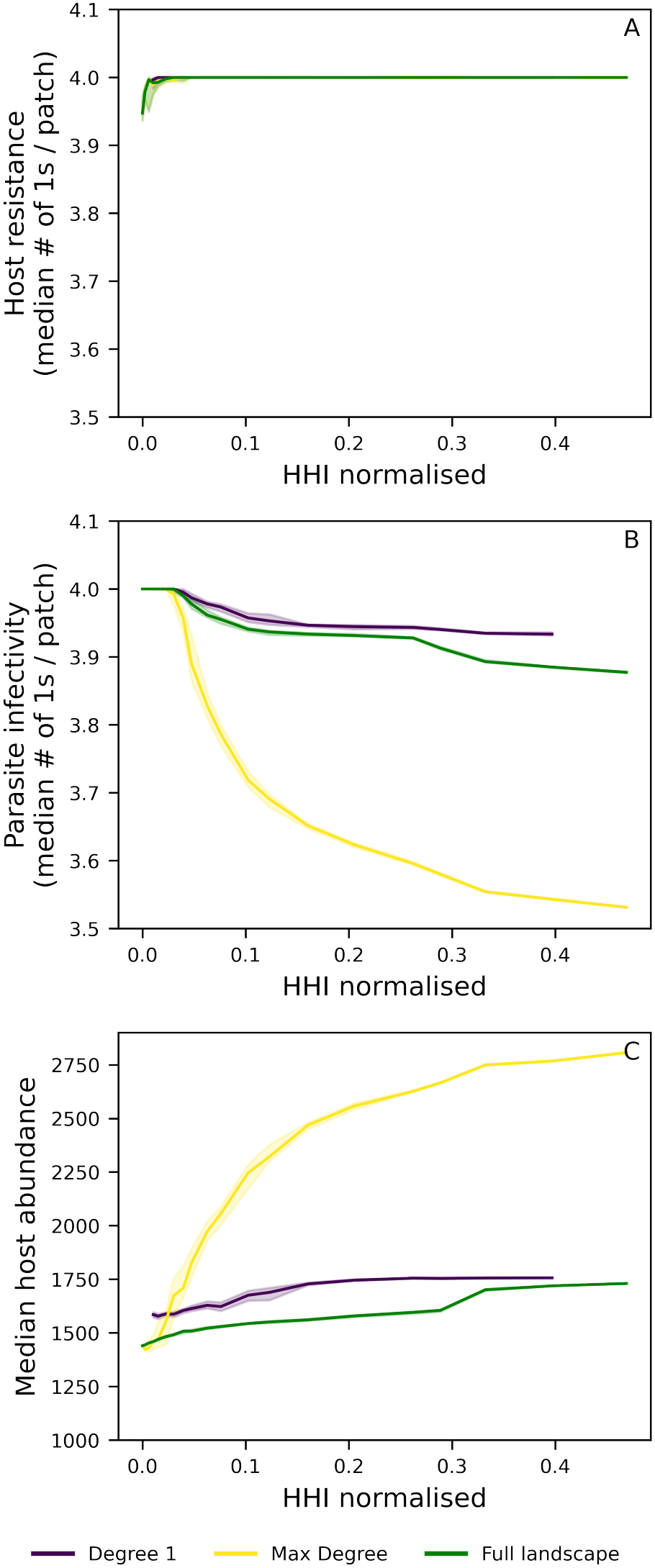
Coevolutionary outcomes and host abundance as a function of degree heterogeneity. The x-axis shows degree heterogeneity measured as the normalised Herfindahl-Hirschman index (*HHI*_norm_; Supplementary Information S4.5.1) across networks generated using edge-rewiring algorithms on a regular 4-connected graph with*n*= 81 patches (see Supplementary Information S4.6) . For each HHI level median and interquartile range (IQR) are computed across networks with different assortativity (quantified by Spearman’s rank correlation coefficient*ρ*; Supplementary Information S4.5.2). Three patch degree subsets are shown:violet-patches with degree 1;yellow-patches with the maximum degree found for each *HHI*_*norm*_ and Spearman’s *ρ*) combination;green-the full network (all patches with degree*≥*1). The three panels correspond to low virulence scaling (*µ*_*V*_ = 0.5) and low transmission cost (*µ*_*T*_ = 0.6) as in Fig. S1 and S29 and marked by the white star in Fig.1. A) Median host resistance (number of “1” alleles per patch). B) Median parasite infectivity (number of “1” alleles per patch). C) Median host abundance per patch. Dispersal is fixed at *d*= 0.3. All other fixed parameters are listed in Table S1. At the bottom, we show a visualisation of networks and corresponding degree-distribution across HHI values.

**Figure S19:**
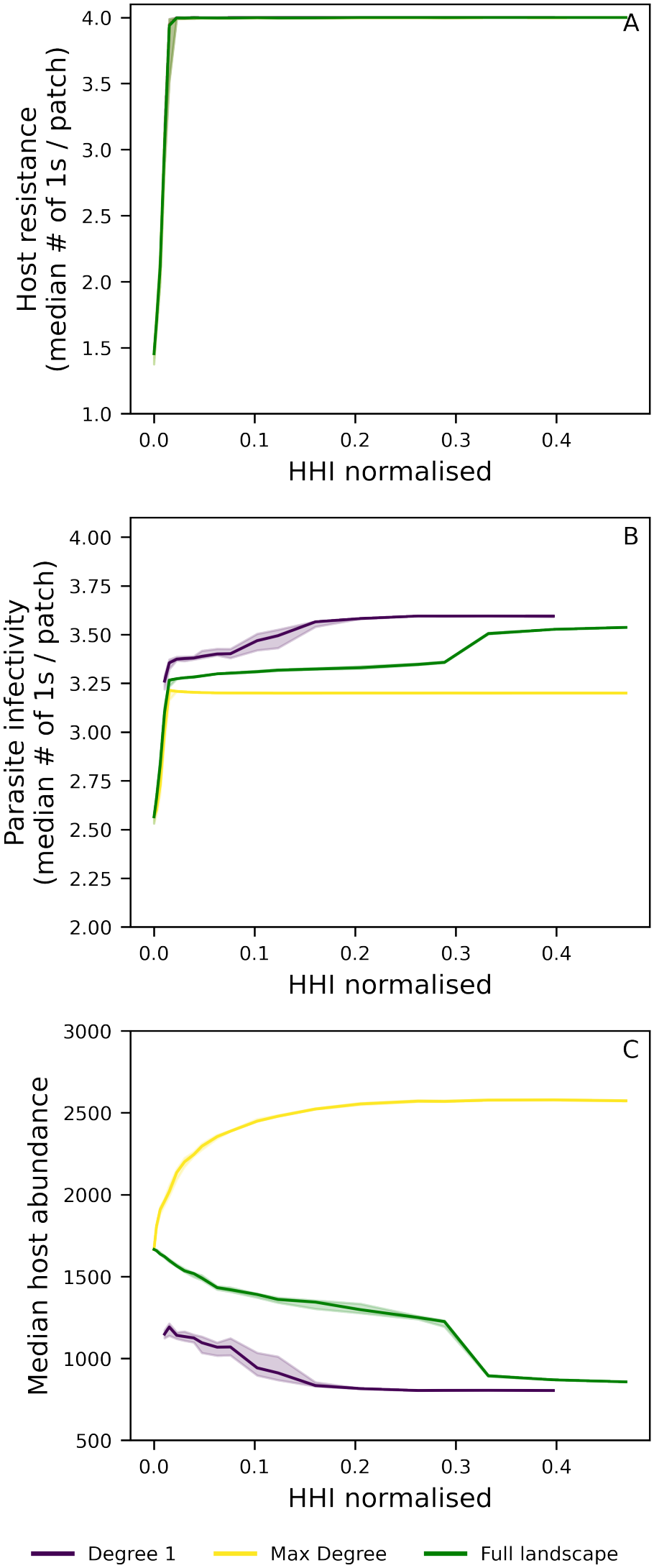
Coevolutionary outcomes and host abundance as a function of degree heterogeneity. The x-axis shows degree heterogeneity measured as the normalised Herfindahl-Hirschman index (*HHI*_norm_; Supplementary Information S4.5.1) across networks generated using edge-rewiring algorithms on a regular 4-connected graph with*n*= 81 patches (see Supplementary Information S4.6). For each HHI level median and interquartile range (IQR) are computed across networks with different assortativity (quantified by Spearman’s rank correlation coefficient*ρ*; Supplementary Information S4.5.2). Three patch degree subsets are shown: violet-patches with degree 1; yellow-patches with the maximum degree found for each *HHI* _*norm*_ and Spearman’s *ρ*) combination; green-the full network (all patches with degree*≥*1). The three panels correspond to low virulence scaling (*µ*_*V*_ = 0.0001) and low transmission cost (*µ*_*T*_ = 0.0001) as in Fig. 2 and S15 and marked by the black star in Fig.1. A) Median host resistance (number of “1” alleles per patch). B) Median parasite infectivity (number of “1” alleles per patch). C) Median host abundance per patch. Dispersal is fixed at *d*= 0.5. All other fixed parameters are listed in Table S1. At the bottom, we show a visualisation of networks and corresponding degree-distribution across HHI values.

**Figure S20:**
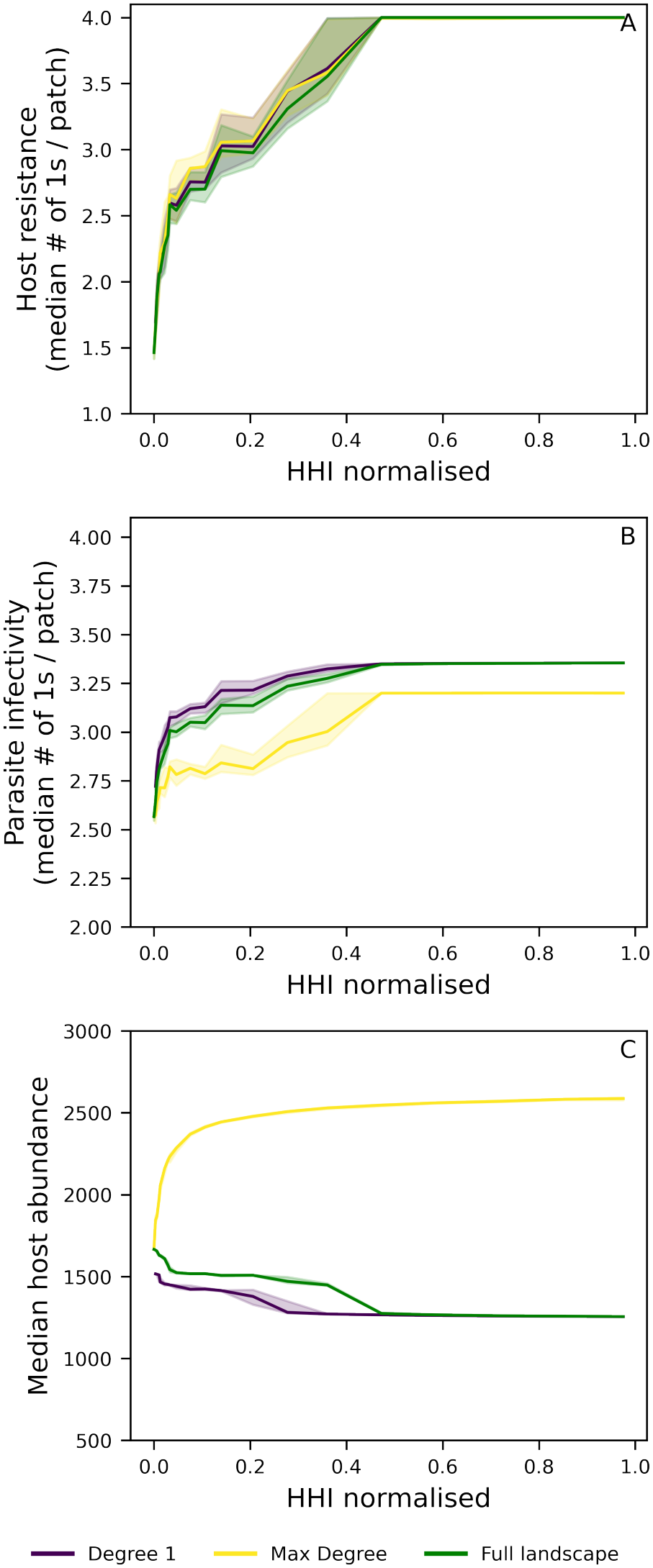
Coevolutionary outcomes and host abundance as a function of degree heterogeneity. The x-axis shows degree heterogeneity measured as the normalised Herfindahl-Hirschman index (*HHI* _*norm*_; Supplementary Information S4.5.1) across networks generated using edge-rewiring algorithms on a regular 2-connected graph with *n* = 81 patches (see Supplementary Information S4.6) . For each HHI level median and interquartile range (IQR) are computed across networks with different assortativity (quantified by Spearman’s rank correlation coefficient *ρ*; Supplementary Information S4.5.2). Three patch degree subsets are shown: violet - patches with degree 1; yellow - patches with the maximum degree found for each *HHI* _*norm*_ and Spearman’s *ρ*) combination; green - the full network (all patches with degree ≥ 1). The three panels correspond to low virulence scaling (*µ*_*V*_ = 0.0001) and low transmission cost (*µ*_*T*_ = 0.0001) as in Fig. 2 and S15 and marked by the black star in Fig.1. A) Median host resistance (number of “1” alleles per patch). B) Median parasite infectivity (number of “1” alleles per patch). C) Median host abundance per patch. Dispersal is fixed at *d* = 0.3. All other fixed parameters are listed in Table S1. At the bottom, we show a visualisation of networks and corresponding degree-distribution across HHI values

**Figure S21:**
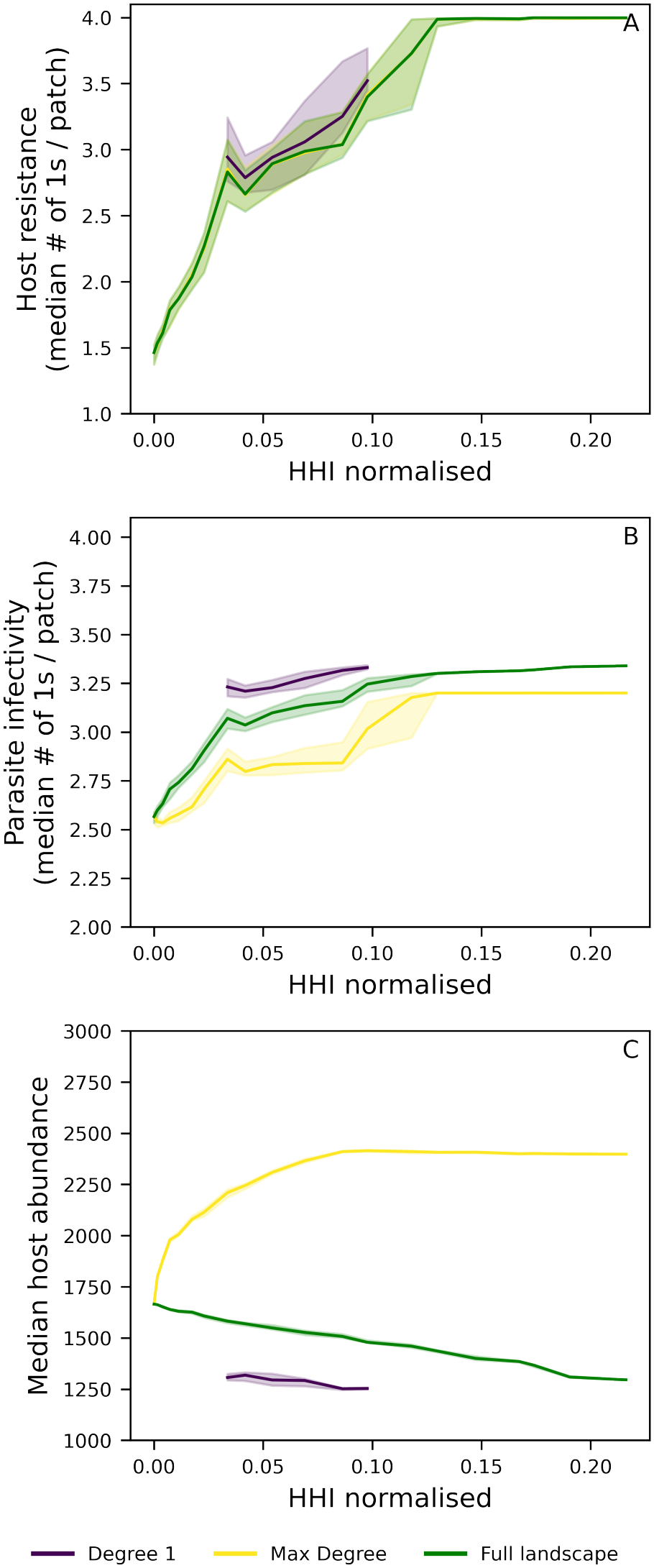
Coevolutionary outcomes and host abundance as a function of degree heterogeneity. The x-axis shows degree heterogeneity measured as the normalised Herfindahl-Hirschman index (*HHI* _*norm*_; Supplementary Information S4.5.1) across networks generated using edge-rewiring algorithms on a regular 8-connected graph with *n* = 81 patches (see Supplementary Information S4.6) . For each HHI level median and interquartile range (IQR) are computed across networks with different assortativity (quantified by Spearman’s rank correlation coefficient *ρ*; Supplementary Information S4.5.2). Three patch degree subsets are shown: violet - patches with degree 1; yellow - patches with the maximum degree found for each *HHI* _*norm*_ and Spearman’s *ρ*) combination; green - the full network (all patches with degree ≥ 1). The three panels correspond to low virulence scaling (*µ*_*V*_ = 0.0001) and low transmission cost (*µ*_*T*_ = 0.0001) as in Fig. 2 and S15 and marked by the black star in Fig.1. A) Median host resistance (number of “1” alleles per patch). B) Median parasite infectivity (number of “1” alleles per patch). C) Median host abundance per patch. Dispersal is fixed at *d* = 0.3. All other fixed parameters are listed in Table S1. At the bottom, we show a visualisation of networks and corresponding degree-distribution across HHI values

**Figure S22:**
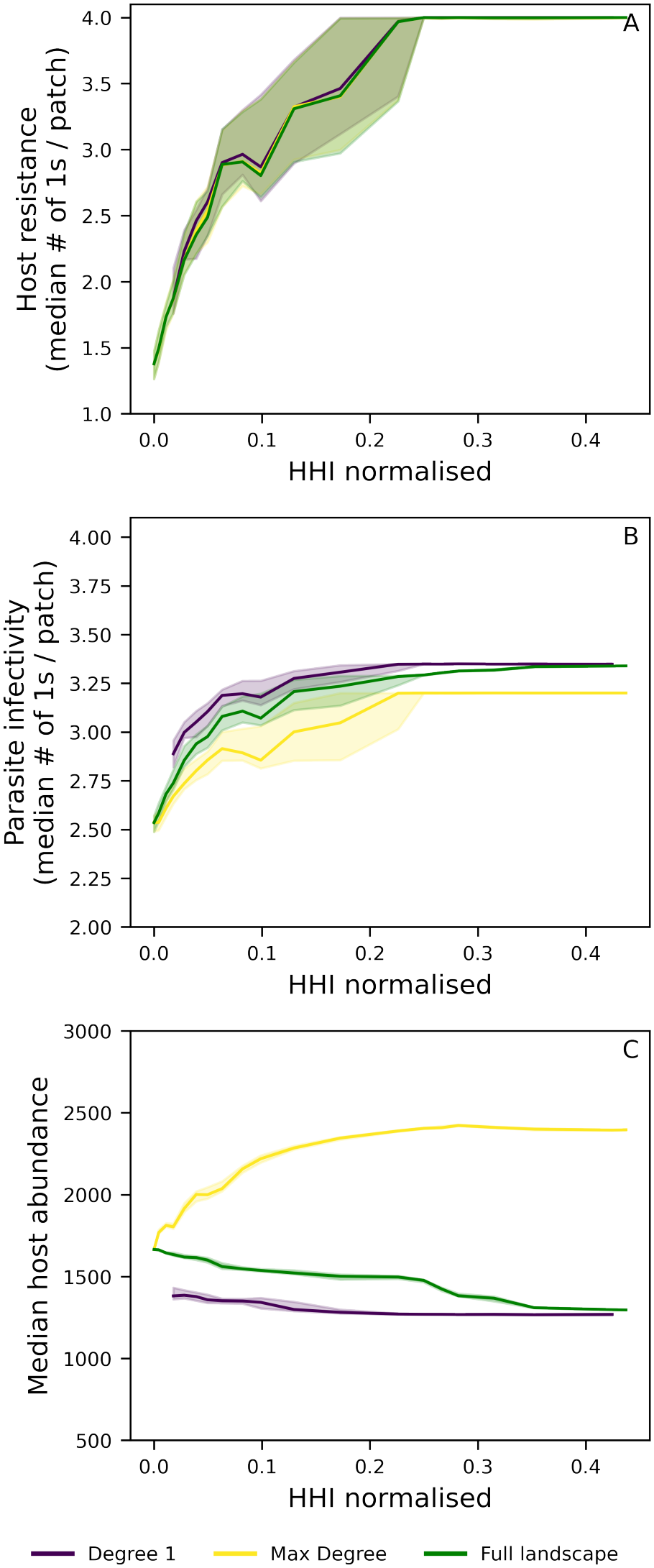
Coevolutionary outcomes and host abundance as a function of degree heterogeneity. The x-axis shows degree heterogeneity measured as the normalised Herfindahl-Hirschman index (*HHI* _*norm*_; Supplementary Information S4.5.1) across networks generated using edge-rewiring algorithms on a regular 4-connected graph with *n* = 40 patches (see Supplementary Information S4.6) . For each HHI level median and interquartile range (IQR) are computed across networks with different assortativity (quantified by Spearman’s rank correlation coefficient *ρ*; Supplementary Information S4.5.2). Three patch degree subsets are shown: violet - patches with degree 1; yellow - patches with the maximum degree found for each *HHI* _*norm*_ and Spearman’s *ρ*) combination; green - the full network (all patches with degree ≥ 1). The three panels correspond to low virulence scaling (*µ*_*V*_ = 0.0001) and low transmission cost (*µ*_*T*_ = 0.0001) as in Fig. 2 and S15 and marked by the black star in Fig.1. A) Median host resistance (number of “1” alleles per patch). B) Median parasite infectivity (number of “1” alleles per patch). C) Median host abundance per patch. Dispersal is fixed at *d* = 0.3. All other fixed parameters are listed in Table S1. At the bottom, we show a visualisation of networks and corresponding degree-distribution across HHI values

**Figure S23:**
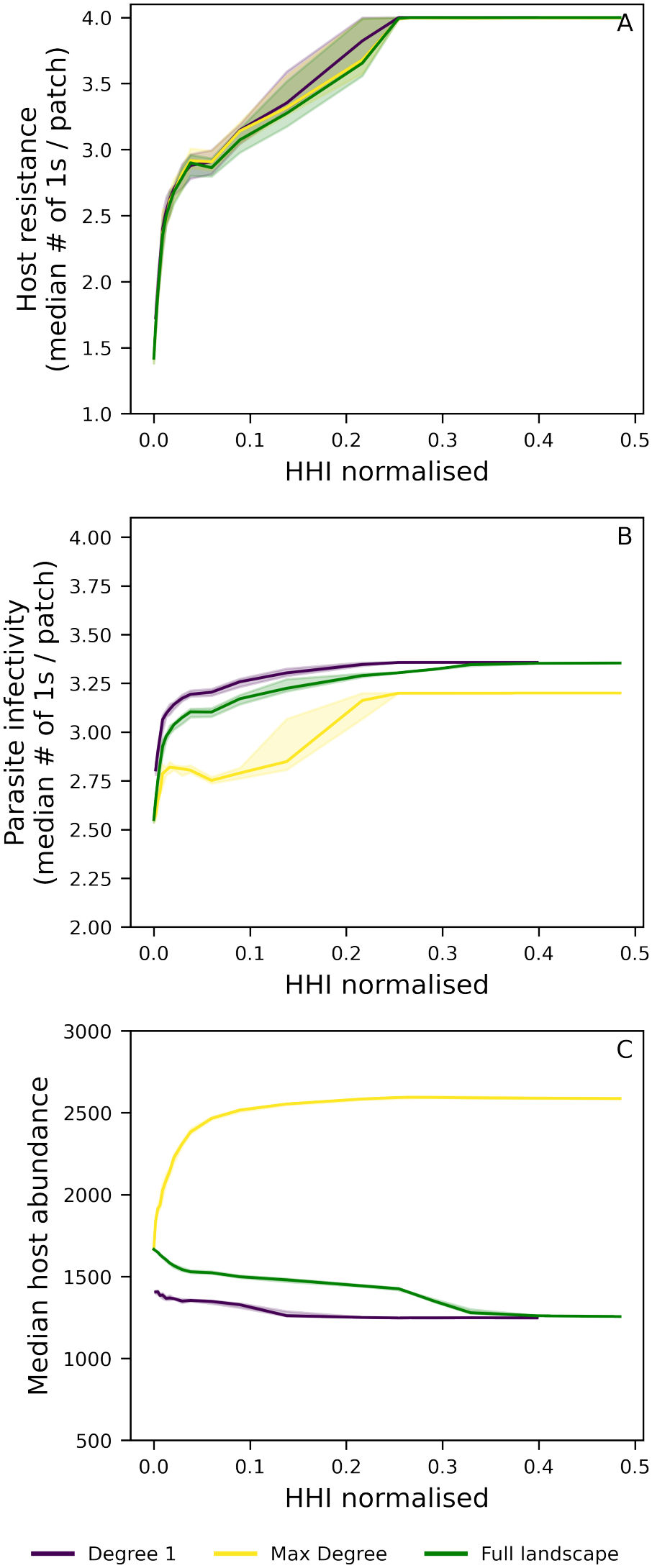
Coevolutionary outcomes and host abundance as a function of degree heterogeneity. The x-axis shows degree heterogeneity measured as the normalised Herfindahl-Hirschman index (*HHI* _norm_; Supplementary Information S4.5.1) across networks generated using edge-rewiring algorithms on a regular 4-connected graph with*n*= 160 patches (see Supplementary Information S4.6) . For each HHI level me- dian and interquartile range (IQR) are computed across networks with different assortativity (quantified by Spearman’s rank correlation coefficient*ρ*; Supplementary Information S4.5.2). Three patch degree subsets are shown:violet- patches with degree 1;yellow- patches with the maximum degree found for each *HHI* _*norm*_ and Spearman’s *ρ*) combination;green- the full network (all patches with degree *≥*1). The three panels correspond to low virulence scaling (*µ* _*V*_ = 0.0001) and low transmission cost (*µ*_*T*_ = 0.0001) as in Fig. 2 and S15 and marked by the black star in Fig.1. A) Median host resistance (number of “1” alleles per patch). B) Median parasite infectivity (number of “1” alleles per patch). C) Median host abundance per patch. Dispersal is fixed at *d*= 0.3. All other fixed parameters are listed in Table S1. At the bottom, we show a visualisation of networks and corresponding degree-distribution

**Figure S24:**
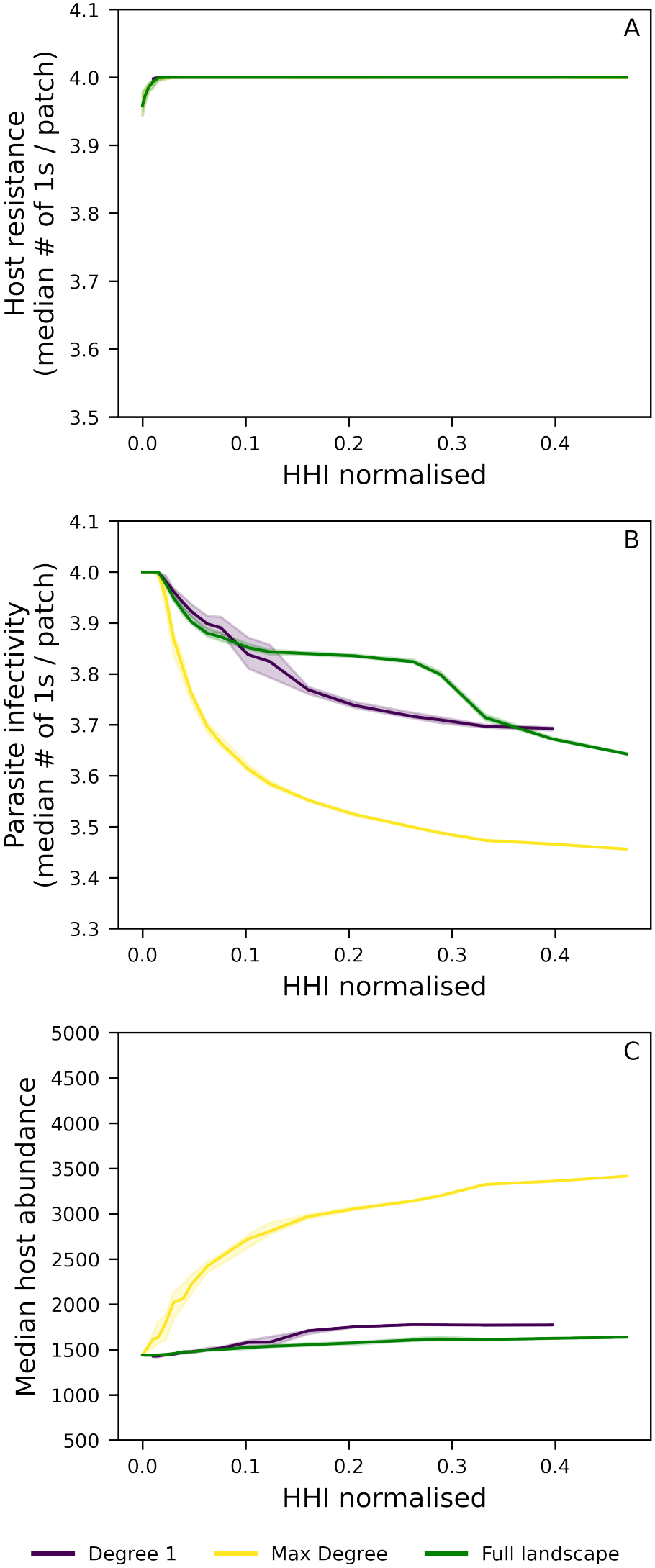
Coevolutionary outcomes and host abundance as a function of degree heterogeneity. The x-axis shows degree heterogeneity measured as the normalised Herfindahl-Hirschman index (*HHI*_norm_; Supplementary Information S4.5.1) across networks generated using edge-rewiring algorithms on a regular 4-connected graph with*n*= 81 patches (see Supplementary Information S4.6) . For each HHI level median and interquartile range (IQR) are computed across networks with different assortativity (quantified by Spearman’s rank correlation coefficient*ρ*; Supplementary Information S4.5.2). Three patch degree subsets are shown:violet- patches with degree 1;yellow- patches with the maximum degree found for each *HHI*_*norm*_ and Spearman’s*ρ*) combination;green- the full network (all patches with degree*≥*1). The three panels correspond to low virulence scaling (*µ*_*V*_ = 0.5) and low transmission cost (*µ*_*T*_ = 0.6) as in Fig. S1 and S29 and marked by the white star in Fig.1. A) Median host resistance (number of “1” alleles per patch). B) Median parasite infectivity (number of “1” alleles per patch). C) Median host abundance per patch. Dispersal is fixed at *d*= 0.5. All other fixed parameters are listed in Table S1. At the bottom, we show a visualisation of networks and corresponding degree-distribution across HHI values.

**Figure S25:**
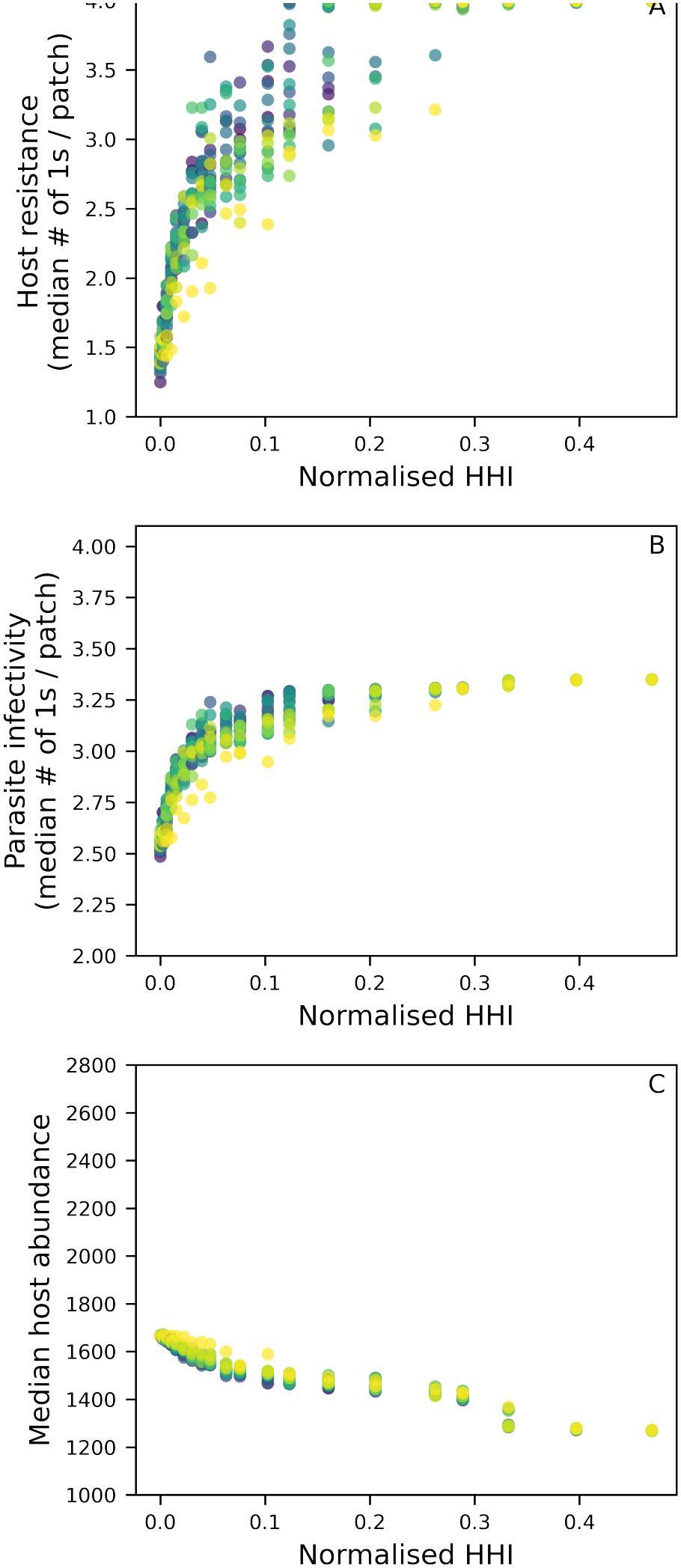
Degree heterogeneity and assortativity 20×20 sweep on a regular 4-connected graph with *n*= 81 patches with dispersal*d*= 0.3. The x-axis shows the normalised Herfindahl-Hirschman index (*HHI*_norm_) values sampled during the sweep (see Supplementary InformationS4.6). For each*HHI* _norm_ level, the median and interquartile range (IQR) of the assortativity parameter are computed across the sweep; assortativity is quantified by Spearman’s *ρ*. Each*HHI* _norm_ value spans 20 assortativity levels, with*ρ*increasing from yellow (low) to blue (high). Panels A, C correspond to low virulence scaling (*µ*_*V*_ = 0.0001) and low transmission cost (*µ*_*T*_ = 0.0001) as in Section 2. Panel A) Median host resistance (number of “1” alleles per patch); Panel B) Median parasite infectivity (number of “1” alleles per patch); Panel C) Median host abundance per patch. Fixed parameters are listed in Table S1.

**Figure S26:**
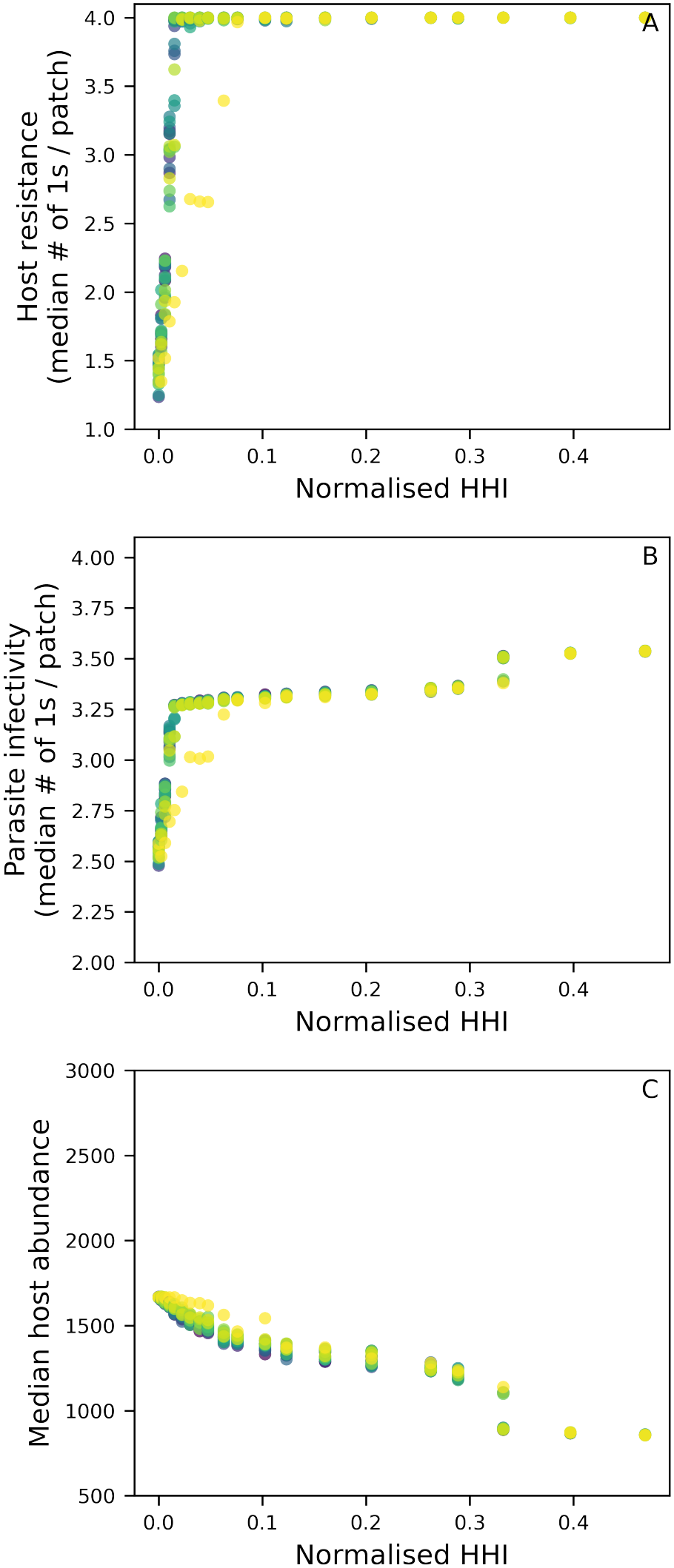
Degree heterogeneity and assortativity 20×20 sweep on a regular 4-connected graph with *n*= 81 patches with dispersal *d*= 0.5. The x-axis shows the normalised Herfindahl-Hirschman index (*HHI*_norm_) values sampled during the sweep (see Supplementary InformationS4.6). For each*HHI* _norm_ level, the median and interquartile range (IQR) of the assortativity parameter are computed across the sweep; assortativity is quantified by Spearman’s *ρ*. Each *HHI* _norm_ value spans 20 assortativity levels, with*ρ*increasing from yellow (low) to blue (high). Panels A, C correspond to low virulence scaling (*µ*_*V*_ = 0.0001) and low transmission cost (*µ*_*T*_ = 0.0001) as in Section 2. Panel A) Median host resistance (number of “1” alleles per patch); Panel B) Median parasite infectivity (number of “1” alleles per patch); Panel C) Median host abundance per patch. Fixed parameters are listed in Table S1.

**Figure S27:**
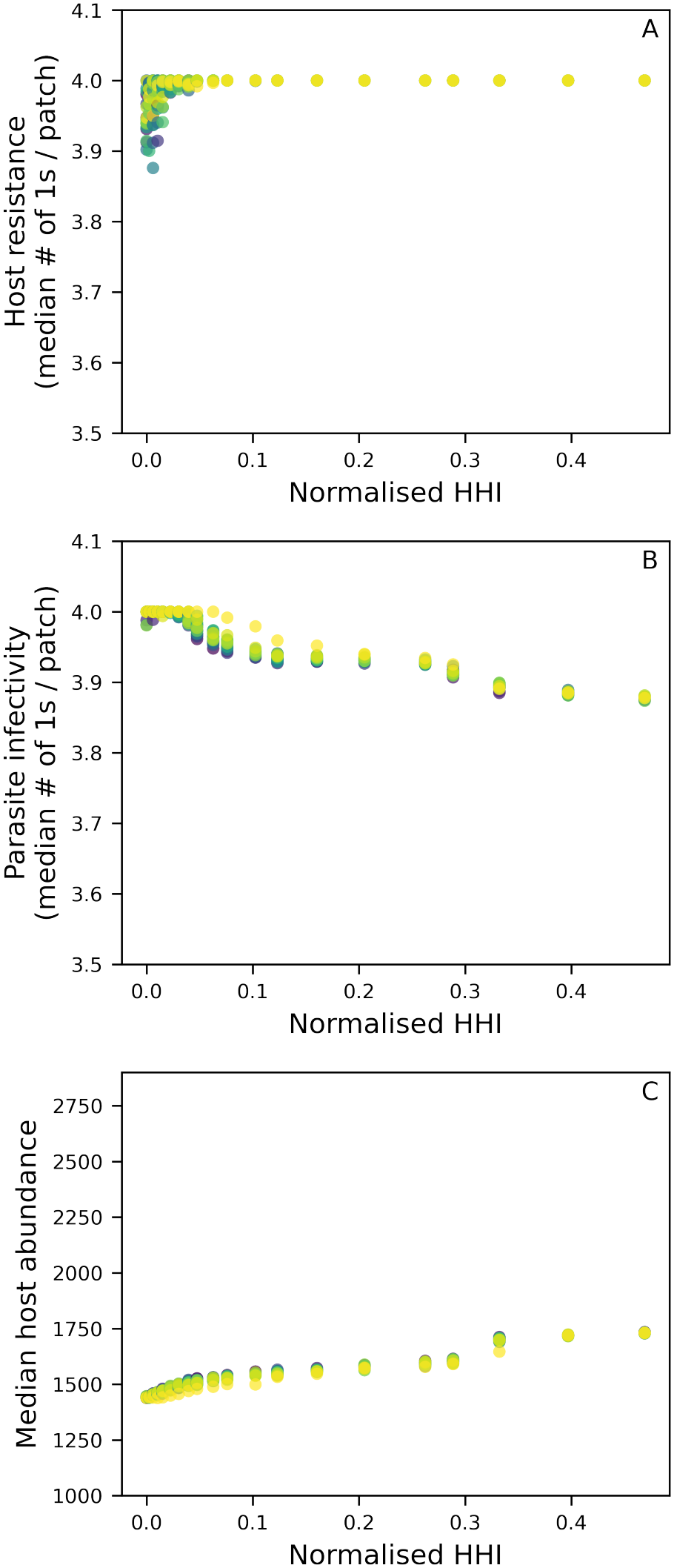
Degree heterogeneity and assortativity 20×20 sweep on a regular 4-connected graph with *n*= 81 patches with dispersal *d*= 0.3. The x-axis shows the normalised Herfindahl-Hirschman index (*HHI*_norm_) values sampled during the sweep (see Supplementary InformationS4.6). For each *HHI* _norm_ level, the median and interquartile range (IQR) of the assortativity parameter are computed across the sweep; assortativity is quantified by Spearman’s *ρ*. Each *HHI* _norm_ value spans 20 assortativity levels, with*ρ*increasing from yellow (low) to blue (high). Panels A, C correspond to low virulence scaling (*µ*_*V*_ = 0.5) and low transmission cost (*µ*_*T*_ = 0.6) as in Section S1. Panel A) Median host resistance (number of “1” alleles per patch); Panel B) Median parasite infectivity (number of “1” alleles per patch); Panel C) Median host abundance per patch. Fixed parameters are listed in Table S1.

**Figure S28:**
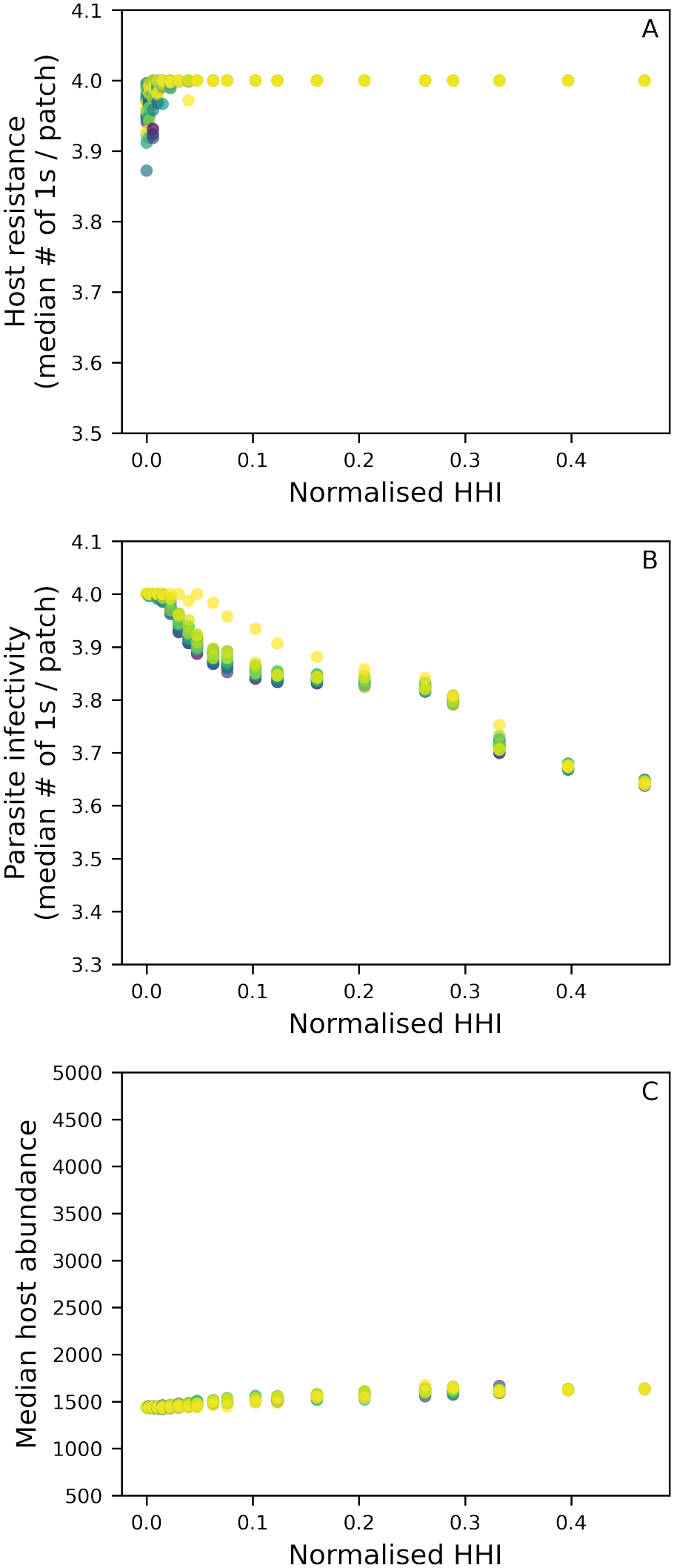
Degree heterogeneity and assortativity 20×20 sweep on a regular 4-connected graph with *n*= 81 patches with dispersal *d*= 0.5. The x-axis shows the normalised Herfindahl-Hirschman index (*HHI*_norm_) values sampled during the sweep (see Supplementary InformationS4.6). For each*HHI* _norm_ level, the median and interquartile range (IQR) of the assortativity parameter are computed across the sweep; assortativity is quantified by Spearman’s*ρ*. Each *HHI* _norm_ value spans 20 assortativity levels, with*ρ*increasing from yellow (low) to blue (high). Panels A, C correspond to low virulence scaling (*µ*_*V*_ = 0.5) and low transmission cost (*µ*_*T*_ = 0.6) as in Section S1. Panel A) Median host resistance (number of “1” alleles per patch); Panel B) Median parasite infectivity (number of “1” alleles per patch); Panel C) Median host abundance per patch. Fixed parameters are listed in Table S1.

**Figure S29:**
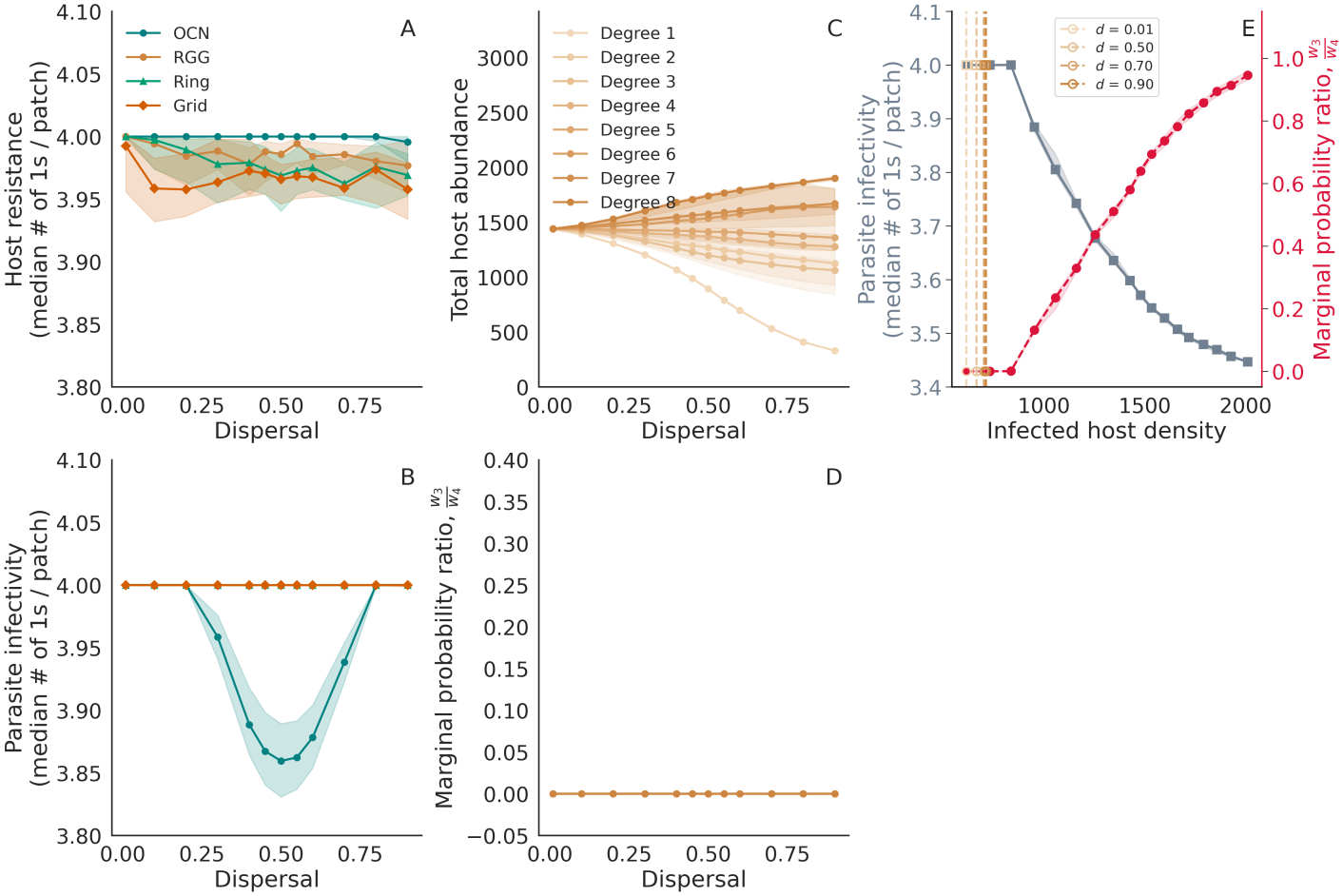
Case study on the impact of dispersal and landscape type for intermediate virulence and transmission costs (*µ*_*V*_ = 0.5,*µ* _*T*_ = 0.6), with a low resistance cost (*µ*_*R*_ = 0.000256). Panel A) Median number of “1” alleles (resistance) in the host population per patch as a function of dispersal. Results are shown for four landscape types: OCN (teal), RGG (orange), Ring (light green), and Grid (dark green). All networks have 81 patches. Shaded regions indicate the interquartile range (IQR) across patches and replicates. Panel B) Median number of “1” alleles (infectivity) in the parasite per patch as a function of dispersal. Shaded regions indicate the IQR across patches and replicates. Panel C) Median host abundance per patch, grouped by patch degree class in the RGG. Shaded regions indicate the IQR across patches and replicates. Panel D) Median ratio of marginal probability of parasite genotypes with 3 “1” or infectivity alleles (*w*_3_) to marginal probability of parasite genotypes with 4 “1” alleles (*w*_4_) as defined by Eqn. S7, computed from the pre-dispersal state and grouped by patch degree in the RGG. Panel E) Parasite infectivity as a function of equilibrium infected host densities in a single patch. The solid grey line shows the median number of “1” infectivity alleles across replicates, and the shaded band gives the IQR. The solid red line shows the median ratio of marginal probability of parasite genotypes with 3 “1” or infectivity alleles (*w*_3_) to marginal probability of parasite genotypes with 4 “1” alleles (*w*_4_) as defined by Eqn. S7 in the single patch. Vertical dashed lines mark the median equilibrium infected host densities in the RGG patches of the target degree found in the simulations at dispersal rate*d*= 0.01,0.5,0.7,0.9, computed from the pre-dispersal state. On each vertical line, the square marker is the median infectivity in those patches (left axis) and the circle marker is the median 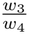 ratio (right axis). Fixed parameters are given in Table S1.

**Figure S30:**
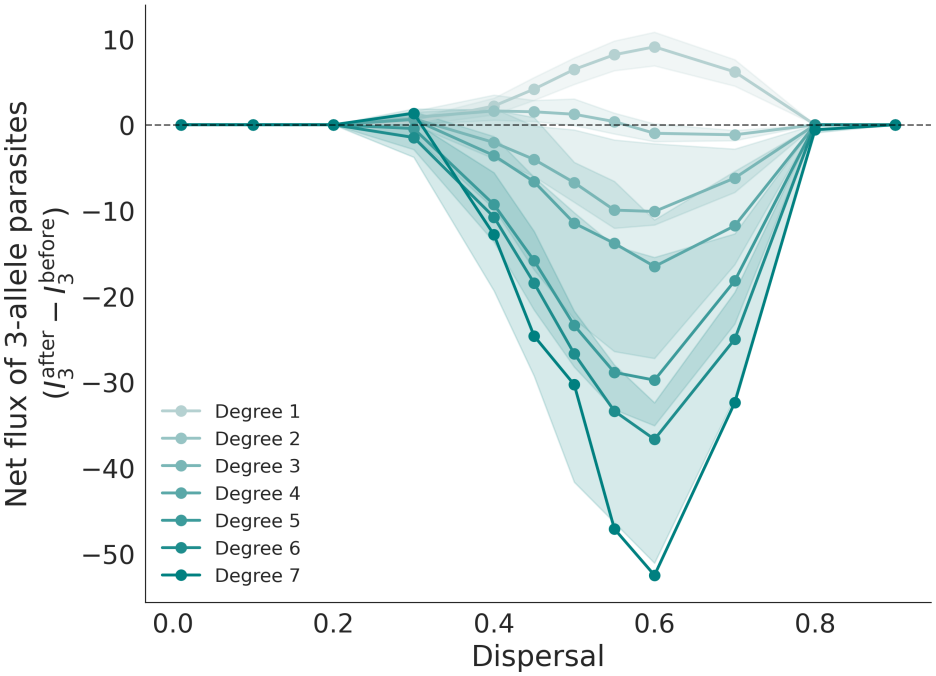
Net flux of parasite sequences with 3 infectivity alleles (1110,1011,1101,1110) in the OCN networks split by degree per patch for*µ* _*V*_ = 0.5,*µ* _*T*_ = 0.6), with a low resistance cost (*µ*_*R*_ = 0.000256) as seen in the analysis of Fig. S1. Flux is measured by subtracting densities of hosts infected by parasites with 3 infectivity alleles after and before dispersal per patch.

### S4 Supplementary Methods

#### S4.1 Empirical grounding for model assumptions

Our model assumes discrete, non-overlapping generations and obligate co-dispersal of parasites with infected hosts. These assumptions are most directly appropriate for host-parasite systems in which transmission is strongly seasonal or pulsed and pathogen movement is coupled to host movement.

The Atlantic salmon–*Gyrodactylus salaris* pathosystem provides a close match to obligate co-dispersal: parasites are transported with migrating fish through river networks (Appleby and Mo, 1997; Jansen and Bakke, 1991). Viral haemorrhagic septicaemia virus in salmonids likewise exhibits temperature-dependent transmission and demography, with transmission driven by seasonal host movement (Kim and Leong, 1999; Garver and Hawley, 2021). Although Atlantic salmon generations overlap, seasonal recruitment produces discrete pulses of susceptible hosts, approximating the discrete-time structure of our model.

Plant pathosystems provide additional examples of strongly seasonal transmission. *Podosphaera plantaginis* is an obligate pathogen of *Plantago lanceolate* that overwinters as resting spores and initiates new epidemics each spring (Jousimo et al., 2014). *Microbotryum* anther-smut fungi are obligate parasites that do not persist in the environment and are transmitted primarily during host flowering (Carlsson-Granér and Thrall, 2015). In both systems, transmission is therefore pulsed and aligned with host phenology.

These plant systems differ from our model in that pathogen dispersal is wind- or vector-mediated rather than strictly co-dispersal with hosts. However, host gene flow through pollen and pathogen movement through spores can be correlated through shared physical processes.

#### S4.2 Full life-cycle algorithm

Each simulation proceeds through the following sequence in every generation:

1. **Dispersal**. Each host disperses with probability *d* from its natal patch to a randomly chosen connected patch. Infected hosts carry their parasites with them.
2. **Reproduction**. Hosts reproduce locally according to Beverton-Holt density dependence (Beverton et al., 1957). In order to model demographic stochasticity, offspring numbers are drawn from a Poisson distribution with expectations given by Eqns. 7 and 8.
3. **Host mutation**. Each locus in each newborn host mutates independently with probability *m* _*H*_ , flipping the allele between 0 and 1.
4. **Transmission**. Newborn susceptible hosts are exposed to the local parasite pool. Infection occurs with probability 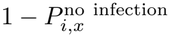, where 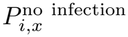 is given by Eqn. 9.
5. **Parasite-genotype selection**. If infection occurs, the establishing parasite genotype is drawn from the local pool with probability proportional to its marginal infection probability.
6. **Parasite mutation**. Each locus in the selected parasite genotype mutates independently with probability *m* _*P*_ .
7. **Generation replacement**. The parental generation is removed, and the newborn generation becomes the next parental generation.

#### S4.3 Transmission formulation

The probability that a newborn host of genotype *i* escapes infection is

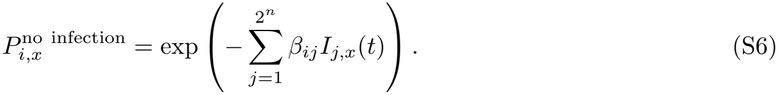

When infection occurs, the probability that parasite genotype *j* establishes the infection is

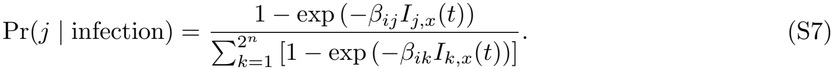

This formulation follows Ashby et al. (2014). It differs from the proportional force-of-infection formulation,

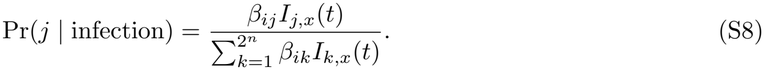

The two formulations converge when infected densities are low. At high infected densities, the marginal-infection formulation saturates: additional parasites of a genotype that already saturates transmission contribute little to infection probability. This gives rarer parasite genotypes a relative advantage and reproduces the empirical observation that per-capita transmission can saturate with infected density (Fenton et al., 2002).

#### S4.4 Landscape construction

##### S4.4.1 Random Geometric Graphs

RGGs were generated in Python using networkx (Hagberg et al., 2008). We placed *n*= 81 points uniformly at random in a unit square and connected pairs of points separated by less than *r*= 0.15. Networks that were not connected were discarded. This procedure yielded a mean degree of approximately 4.5. We generated 20 independent stochastic realisations.

##### S4.4.2 Optimal Channel Networks

OCNs were generated on an un-aggregated grid using code from Deshpande et al. (2025), implementing the approach of Carraro et al. (2020), with a fixed outlet. We used *n*= 81 patches, yielding a mean degree of approximately 2. We used unweighted, undirected graphs; explicit riverine directionality would likely strengthen the ecological asymmetries already captured by degree heterogeneity (Fronhofer and Altermatt, 2017). We generated 20 independent stochastic realisations.

##### S4.4.3 Homogeneous controls

Because OCNs and RGGs differ substantially in mean degree (see above: 4.5 for RGGs and 2 for OCNs), we included homogeneous controls for mean degree. A regular grid with periodic boundary conditions and a connectivity of 4 served as the homogeneous counterpart to the RGG, and a ring served as the homogeneous counterpart to the OCN.

#### S4.5 Network Metrics

##### S4.5.1 Degree-heterogeneity (HHI)

We quantified degree-heterogeneity using the normalised Herfindahl-Hirschman Index (Hirschman, 1980).

Let *d* _*i*_ be the degree of node *i*, and let 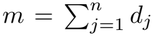 be the total degree sum then raw concentration index is:

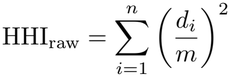

To make values comparable across networks of different sizes, we normalised HHI _raw_ against its theoretical minimum and maximum (Estrada, 2010). The minimum is 1*/n*, achieved when all degrees are equal (a regular graph). The maximum is *n/*(4(*n−*1)), which corresponds to a perfect star graph (one node of degree *n−*1, all others degree 1). Thus:

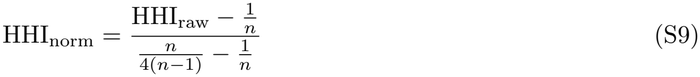

This yields a metric ranging from 0 to 1. It is important to note that the theoretical maximum is not always reachable: when the total number of edges is more than*n−*1, a pure star is structurally impossible while keeping number of edges equal. But HHI still scales relative to this upper bound, maintaining a metric that allows direct comparison across networks with different number of nodes.

##### S4.5.2 Assortativity (Spearman’s*ρ*)

To quantify the tendency of nodes to connect to others of similar degree, we use Spearman’s *ρ*(Litvak and Van Der Hofstad, 2013) of the degrees at the two endpoints of each edge. For every edge (*u, v*) in the graph, we form the pair (*d*_*u*_, *d*_*v*_), where *d* _*u*_ is the degree of node*u*. The Spearman’s*ρ*is computed as the rank correlation between the two sequences*{d* _*u*_*}*and*{d* _*v*_*}*over all edges:

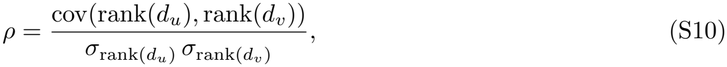

where ranks are computed separately for the two degree lists. A positive*ρ*indicates assortative mixing (hubs tend to connect to hubs), while a negative *ρ* indicates disassortative mixing (hubs connect to leaves). This metric is not fixed by the degree sequence, it depends on how the edges are arranged, making it the ideal complement to HHI, which only measures degree inequality. For a given degree sequence, different rewiring patterns can yield a range of *ρ* values, from highly assortative to highly disassortative.

#### S4.6 HHI and Spearman’s*ρ*nested exploration algorithm

##### S4.6.1 HHI sweep

Starting from a user-specified regular graph, we used a hub-targeted rewiring algorithm to maximise degree heterogeneity while preserving the total number of edges and connectivity. At each step, a hub node was selected with probability proportional to its current degree raised to a power 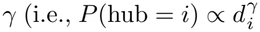, with*γ*= 1.5). An edge not incident to the hub was then randomly chosen and rewired to connect the hub to a randomly selected non-neighbour. This operation concentrated degree into the hub while reducing the degrees of the two endpoints of the removed edge. Moves were accepted only if they increased the normalised HHI and maintained graph connectivity. The algorithm ran for a maximum of 200,000 steps, and graphs were recorded whenever HHI reached a new all-time high, ensuring a strictly monotonic trajectory of increasing degree heterogeneity.

##### S4.6.2 Nested Spearman’s *ρ* exploration for each fixed degree sequence

For each HHI level sampled from the trajectory (i.e., for each fixed degree sequence), we explore the full range of possible Spearman’s *ρ* values using three complementary strategies based on degree-preserving double-edge swaps. All swaps maintain the total number of edges and graph connectivity (checked via bridge tests), and they preserve the degree sequence exactly, so HHI remains constant within each level.

1. Random walk: We perform an unbiased random walk for 200,000 steps, recording a snapshot every 100 steps and additionally recording the graph whenever a new global minimum or maximum *ρ* is encountered (checked every 100 steps). This provides a broad, unbiased sample of the*ρ*landscape.
2. Greedy maximisation: We run a separate greedy search for up to 5,000 steps, accepting only swaps that strictly increase *ρ*. This drives the graph towards assortative hub-hub connections, yielding the upper bound of *ρ* for that degree sequence.
3. Greedy minimisation: We run a separate greedy search for up to 5,000 steps, accepting only swaps that strictly decrease *ρ*. This drives the graph towards disassortative hub-leaf connections, yielding the lower bound of *ρ* for that degree sequence.

## References

Agrawal, A. and Lively, C. 2003. Modelling infection as a two-step process combining gene-for-gene and matching-allele genetics. – Proceedings of the Royal Society B: Biological Sciences 270(1512): 323–334.

Alizon, S. and van Baalen, M. 2005. Emergence of a convex trade-off between transmission and virulence. – The American Naturalist 165(6): E155–E167.

Anderson, R. M. and May, R. M. 1982. Coevolution of hosts and parasites. – Parasitology 85(2): 411–426.

Appleby, C. and Mo, T. A. 1997. Population dynamics of Gyrodactylus salaris (Monogenea) infecting Atlantic salmon, Salmo salar, parr in the River Batnfjordselva, Norway. – The Journal of Parasitology 83(1): 23–30.

Ashby, B., Gupta, S. and Buckling, A. 2014. Spatial structure mitigates fitness costs in host-parasite coevolution. – The American Naturalist 183(3): E64–E74.

Ashby, B., Iritani, R., Best, A., White, A. and Boots, M. 2019. Understanding the role of eco-evolutionary feedbacks in host-parasite coevolution. – Journal of Theoretical Biology 464:115–125.

Bailey, V. A. 1931. The interaction between hosts and parasites. – The Quarterly Journal of Mathematics os-2(1): 68–77.

Beverton, R. J. H., Ministry of Agriculture, Fisheries and Food and Holt, S. J. 1957. On the dynamics of exploited fish populations. – HMSO.

Boots, M. and Sasaki, A. 1999. Small worlds and the evolution of virulence: infection occurs locally and at a distance. – Proceedings of the Royal Society B: Biological Sciences 266(1432): 1933–1938.

Boussange, V. and Pellissier, L. 2022. Eco-evolutionary model on spatial graphs reveals how habitat structure affects phenotypic differentiation. – Communications Biology 5(1): 668.

Buckingham, L. J. and Ashby, B. 2022. Coevolutionary theory of hosts and parasites. – Journal of Evolutionary Biology 35(2): 205–224.

Bullock, J. M., Bonte, D., Pufal, G., da Silva Carvalho, C., Chapman, D. S., García, C., García, D., Matthysen, E. and Delgado, M. M. 2018. Human-mediated dispersal and the rewiring of spatial networks. – Trends in Ecology & Evolution 33(12): 958–970.

Carlsson-Granér, U. and Thrall, P. H. 2015. Host resistance and pathogen infectivity in host populations with varying connectivity. – Evolution 69(4): 926–938.

Carraro, L., Bertuzzo, E., Fronhofer, E. A., Furrer, R., Gounand, I., Rinaldo, A. and Altermatt, F. 2020. Generation and application of river network analogues for use in ecology and evolution. – Ecology and Evolution 10(14): 7537–7550.

Colizza, V. and Vespignani, A. 2007. Invasion threshold in heterogeneous metapopulation networks. – Physical Review Letters 99(14): 148701.

Deshpande, J. N., Dakos, V., Kaltz, O. and Fronhofer, E. A. 2025a. Landscape structure as a driver of eco-evolution in host–parasite systems. – Evolution Letters 9(3): 367–378.

Deshpande, J. N., Pallagatti, R. S., Dakos, V., Kaltz, O. and Fronhofer, E. A. 2025b. Co-evolution of host dispersal and parasite virulence in complex landscapes. – Evolution 80(2): 487–495.

Fenton, A. and Brockhurst, M. A. 2007. Epistatic interactions alter dynamics of multilocus gene-for-gene coevolution. – PLoS One 2(11): e1156.

Flor, H. H. 1956. The complementary genic systems in flax and flax rust. – Advances in genetics 8:29–54.

Fronhofer, E. A. and Altermatt, F. 2017. Classical metapopulation dynamics and eco-evolutionary feed-backs in dendritic networks. – Ecography 40(12): 1455–1466.

Gandon, S. 2002. Local adaptation and the geometry of host-parasite coevolution. – Ecology Letters 5(2): 246–256.

Gandon, S., van Baalen, M. and Jansen, V. A. A. 2002. The evolution of parasite virulence, superinfection, and host resistance. – The American Naturalist 159(6): 658–669.

Garver, K. and Hawley, L. 2021. Characterization of viral haemorrhagic septicaemia virus (VHSV) to inform pathogen transfer risk assessments in British Columbia. – Canadian Science Advisory Secretariat.

Gilarranz, L. J. and Bascompte, J. 2012. Spatial network structure and metapopulation persistence. – Journal of Theoretical Biology 297:11–16.

Guivier, E., Galan, M., Chaval, Y., Xuéreb, A., Ribas Salvador, A., Poulle, M.-L., Voutilainen, L., Henttonen, H., Charbonnel, N. and Cosson, J.-F. 2011. Landscape genetics highlights the role of bank vole metapopulation dynamics in the epidemiology of Puumala hantavirus. – Molecular Ecology 20(17): 3569–3583.

Gustafsson, L. and Sternad, M. 2007. Bringing consistency to simulation of population models–Poisson Simulation as a bridge between micro and macro simulation. – Mathematical Biosciences 209(2): 361–385.

Hassell, M. P. 2000. Host-parasitoid population dynamics. – Journal of Animal Ecology 69(4): 543–566.

Höckerstedt, L., Numminen, E., Ashby, B., Boots, M., Norberg, A. and Laine, A.-L. 2022. Spatially structured eco-evolutionary dynamics in a host-pathogen interaction render isolated populations vulnerable to disease. – Nature Communications 13(1): 6018.

Jousimo, J., Tack, A. J., Ovaskainen, O., Mononen, T., Susi, H., Tollenaere, C. and Laine, A.-L. 2014. Ecological and evolutionary effects of fragmentation on infectious disease dynamics. – Science 344(6189): 1289–1293.

Kermack, W. O. and McKendrick, A. G. 1927. A contribution to the mathematical theory of epidemics. – Proceedings of the Royal Society of London. Series A, Containing Papers of a Mathematical and Physical Character 115(772): 700–721.

Kim, Y. S. and Ashby, B. 2026. Coevolutionary cycling in allele frequencies and the evolution of virulence. – Evolution 80(1): 282–290.

Kininmonth, S., Drechsler, M., Johst, K. and Possingham, H. P. 2010. Metapopulation mean life time within complex networks. – Marine Ecology Progress Series 417:139–149.

Lion, S. and Boots, M. 2010. Are parasites “prudent” in space?. – Ecology Letters 13(10): 1245–1255.

Litvak, N. and Van Der Hofstad, R. 2013. Uncovering disassortativity in large scale-free networks. – Physical Review E-Statistical, Nonlinear, and Soft Matter Physics 87(2): 022801.

North, A. R. and Godfray, H. C. J. 2017. The dynamics of disease in a metapopulation: The role of dispersal range. – Journal of Theoretical Biology 418:57–65.

Parratt, S. R., Numminen, E. and Laine, A.-L. 2016. Infectious disease dynamics in heterogeneous landscapes. – Annual Review of Ecology, Evolution, and Systematics 47(1): 283–306.

Peters, M. A. E., Mideo, N. and MacPherson, A. 2023. The maintenance of genetic diversity under host-parasite coevolution in finite, structured populations. – Journal of Evolutionary Biology 36(9): 1328–1341.

Rinaldo, A., Rigon, R., Banavar, J. R., Maritan, A. and Rodriguez-Iturbe, I. 2014. Evolution and selection of river networks: Statics, dynamics, and complexity. – Proceedings of the National Academy of Sciences 111(7): 2417–2424.

Sasaki, A. 2000. Host-parasite coevolution in a multilocus gene-for-gene system. – Proceedings of the Royal Society B: Biological Sciences 267(1458): 2183–2188.

Sasaki, A., Hamilton, W. D. and Úbeda, F. 2002. Clone mixtures and a pacemaker: new facets of Red-Queen theory and ecology. – Proceedings of the Royal Society B: Biological Sciences 269(1493): 761–772.

Sato, M., Dieckmann, U. and Sasaki, A. 2024. Metapopulation heterogeneities in host mobility, productivity, and immunocompetency always increase virulence and infectiousness. – Proceedings of the National Academy of Sciences 121(52): e2309272121.

Tadiri, C. P., Scott, M. E. and Fussmann, G. F. 2018. Microparasite dispersal in metapopulations: a boon or bane to the host population?. – Proceedings of the Royal Society B: Biological Sciences 285(1885): 20181519.

Testa, S., Berger, S., Piccardi, P., Oechslin, F., Resch, G. and Mitri, S. 2019. Spatial structure affects phage efficacy in infecting dual-strain biofilms of Pseudomonas aeruginosa. – Communications Biology 2(1): 405.

Thompson, J. N. 2019. The geographic mosaic of coevolution. – University of Chicago Press.

Tian, H., Liu, Y., Li, Y., Wu, C.-H., Chen, B., Kraemer, M. U. G., Li, B., Cai, J., Xu, B., Yang, Q., Wang, B., Yang, P., Cui, Y., Song, Y., Zheng, P., Wang, Q., Bjornstad, O. N., Yang, R., Grenfell, B. T., Pybus, O. G. and Dye, C. 2020. An investigation of transmission control measures during the first 50 days of the COVID-19 epidemic in China. – Science 368(6491): 638–642.

Volz, E. M., Miller, J. C., Galvani, A. and Meyers, L. A. 2011. Effects of Heterogeneous and Clustered Contact Patterns on Infectious Disease Dynamics. – PLOS Computational Biology 7(6): e1002042.

Walsman, J. C., Duffy, M. A., Cáceres, C. E. and Hall, S. R. 2023. Resistance is futile: Weaker selection for resistance by abundant parasites increases prevalence and depresses host density. – The American Naturalist 201(6): 864–879.

## References

Deshpande, J. N., Pallagatti, R. S., Dakos, V., Kaltz, O. and Fronhofer, E. A. 2025. Co-evolution of host dispersal and parasite virulence in complex landscapes. – Evolution 80(2): 487–495.

Estrada, E. 2010. Quantifying network heterogeneity. – Physical Review E-Statistical, Nonlinear, and Soft Matter Physics 82(6): 066102.

Fenton, A., Fairbairn, J. P., Norman, R. and Hudson, P. J. 2002. Parasite transmission: reconciling theory and reality. – Journal of Animal Ecology 71(5): 893–905.

Hagberg, A., Swart, P. J. and Schult, D. A. 2008. Exploring network structure, dynamics, and function using NetworkX. – Tech. rep., Los Alamos National Laboratory (LANL).

Hirschman, A. O. 1980. National power and the structure of foreign trade. – vol. 105. Univ of California Press.

Jansen, P. and Bakke, T. 1991. Temperature-dependent reproduction and survival of Gyrodactylus salaris Malmberg, 1957 (Platyhelminthes: Monogenea) on Atlantic salmon (Salmo salar L.). – Parasitology 102(1): 105–112.

Kim, C. H. and Leong, J.-A. 1999. FISH VIRUSES. – In: Granoff, A.and Webster, R. G. (eds.), Encyclopedia of Virology (Second Edition). Elsevier, pp. 558–568. Second edition ed.

Lion, S. and Metz, J. A. 2018. Beyond R0 maximisation: on pathogen evolution and environmental dimensions. – Trends in Ecology & Evolution 33(6): 458–473.

